# A Multiomics, Spatiotemporal, and Single Cell Atlas for Mapping Cell-Type-Specific Dysregulation at the Maternal-Fetal Interface

**DOI:** 10.1101/2024.01.18.576324

**Authors:** Cheng Wang, Yan Zhou, Yuejun Jessie Wang, Tuhin Kumar Guha, Zhida Luo, Tara I. McIntyre, Marisa E. Schwab, Brittany R. Davidson, Gabriella C. Reeder, Ronald J. Wong, Sarah England, Juan M. Gonzalez, Robert Blelloch, Alexis J. Combes, Linda C. Giudice, Adrian Erlebacher, Tippi C. MacKenzie, David K. Stevenson, Gary M. Shaw, Michael P. Snyder, Susan J. Fisher, Virginia D. Winn, Jingjing Li

## Abstract

The placenta, the first organ to functionally mature, undergoes disordered development in many pregnancy complications. Molecular investigations have been hampered by the extreme cellular heterogeneity of the placenta, and this complexity is further exaggerated at the maternal-fetal interface where maternal and fetal cells co-mingle. We generated the paired single nucleus epigenomes and transcriptome for each of ∼200,000 cells at the human maternal-fetal interface from early pregnancy to term. These data identified cell-type-specific transcriptional regulatory programs and uncovered key transcription factors driving the lineage differentiation of placental cytotrophoblasts. Integrating spatial single cell proteomics profiling, we localized the observed cell types *in situ*, and characterized the dynamic stages and distinct features of endothelial cells of maternal spiral arteries remodeled by extravillous cytotrophoblasts. Integrative analyses of the single cell data across gestation enabled fine-mapping of the developmental trajectories of cytotrophoblasts and decidual stromal cells, and defining the signature molecular profiles of known and novel cell (sub)types. To demonstrate clinical value, we integrated the reference single cell data with large-scale population genomes from pregnancy complications and identified the most vulnerable maternal and fetal cell types in preeclampsia, preterm birth, and miscarriage. This study presents the most comprehensive placental and decidual single cell resource across gestation to date, reveals new insights into the drivers of normal human placentation, and uncovers the cellular basis of dysfunction associated with common pregnancy complications.

## Introduction

Pregnancy-associated complications threaten maternal and fetal health. For example, miscarriage occurs in 15% of recognized pregnancies(Quenby et al., 2021). Worldwide, preeclampsia is associated with over 70,000 maternal and 500,000 fetal deaths annually(Rana et al., 2019). Preterm birth is the leading contributor to neonatal death affecting 10% of pregnancies(Lee et al., 2019). Many pregnancy complications are associated with dysregulated placental development, but the molecular etiologies have largely remained unclear. The placenta is a transient organ, whose morphology and function are highly dynamic over pregnancy. At implantation, specialized placental cells, termed cytotrophoblasts (CTBs), burrow into the uterus, initiating attachment. Chorionic villi cytotrophoblast (VCT) progenitors either fuse to form syncytiotrophoblasts (SCTs), a transport epithelium, or give rise to extravillous trophoblast cells (EVTs) that invade the decidua and remodel the spiral arteries (SAs). By the end of the first trimester, the latter process redirects uterine blood flow to the intervillous space, supporting rapid growth of the placenta and the embryo/fetus. The cells’ dynamic roles at the maternal-fetal interface require coordinated epigenetic and transcriptional regulation to tune responses to the different physiological states and cellular environments encountered over gestation. It is reasonable to expect that dysregulation at any level could compromise pregnancy outcomes.

Molecular characterization of placental development, in particular the portion that interfaces with the uterus, has been challenging owing to the unique mosaic composition of this interface where maternal and fetal cells co-mingle. As such, understanding molecular dynamics of placental development requires resolution at a single cell level. However, technological advancements have only recently made the required resolution possible. To date, published single cell RNA-seq analyses of the human placenta and the maternal-fetal interface targeted specific developmental stages (details are summarized in **Table S1**)(Arutyunyan et al., 2023; Liu et al., 2018; Marsh et al., 2022; Pique-Regi et al., 2019; Suryawanshi et al., 2018; Tsang et al., 2017; Vento-Tormo et al., 2018). Thus, the developmental trajectories of cells over gestation have not been fully characterized. For example, single cell RNA-seq limited to the first trimester captured very few endovascular EVTs(Arutyunyan et al., 2023; Vento-Tormo et al., 2018), which remodel maternal SAs, a process that is most active at mid-gestation. This likely explains why the number of endovascular EVTs did not proportionally grow with substantially increased numbers of sequenced early gestation cells(Arutyunyan et al., 2023). Likewise, analyzing term samples omits key developmental events in early gestation. Unlike single cell transcriptome profiling, imaging-based cell characterization using a small protein panel (∼30 proteins enriched for immune functions) would not allow agnostic discovery of new cell (sub)types at a global level(Greenbaum et al., 2023; Keren et al., 2019). While the existing single cell data characterizing the maternal-fetal interface are broadly confirmatory of cellular heterogeneity at a given developmental stage, it remains unknown whether there are major developmental trajectories across gestation on which seemingly heterogeneous cell states could converge. Further, translational insights into cell-type-specific contribution to pregnancy complications have not yet been derived.

Significantly extending the existing single cell transcriptomic analysis, we performed single cell multiomics profiling across gestation of the maternal-fetal interface in normal pregnancies, generating the paired epigenome (ATAC-seq) and transcriptome data (RNA-seq) for each of ∼200,000 cells/nuclei from gestational week (GW) 5 to GW 39 (**Fig.1A**). Our post-QC high-quality multiomics_data profiled on average >8,000 cells/nuclei per sample, which nearly doubled or tripled the per sample throughput of most previous studies (**Table S1**). The higher level of cell abundance and the inclusion of key developmental stages across gestation led us to identify novel cell types and enabled fine mapping of cell developmental trajectories across gestation. Compared with existing placental and decidual single cell data primarily based on RNA-seq analyses (**Table S1**), our paired epigenome and transcriptome for each cell demonstrated enhanced power in reconstructing cell-type-specific transcriptional regulatory networks. Further integration with spatial single cell proteomics profiling precisely localized the observed cell types at the maternal-fetal interface *in situ*. Our single cell multiomics data integration enabled our discovery of distinct endothelial states in response to EVT remodeling of maternal SAs, the reconstruction of complete cell developmental trajectories at the maternal fetal interface, and the identification of novel trophoblast and decidual stromal cell (sub)types. To demonstrate translational utility, we integrated the single-cell multiomics data with large-scale maternal and fetal genomes from pregnancies complicated by preeclampsia, preterm birth, or miscarriage. This novel analysis identified vulnerable maternal and/or fetal cell types specific to each condition, suggesting molecular etiologies and potential targets for future therapeutic development. Overall, our study provides new insights into the development of the maternal-fetal interface and unraveled key contributory cell types in major pregnancy complications. We provide a UCSF Single Cell Data Browser to share this data resources with the community (see Data Availability section).

**Fig. 1.**
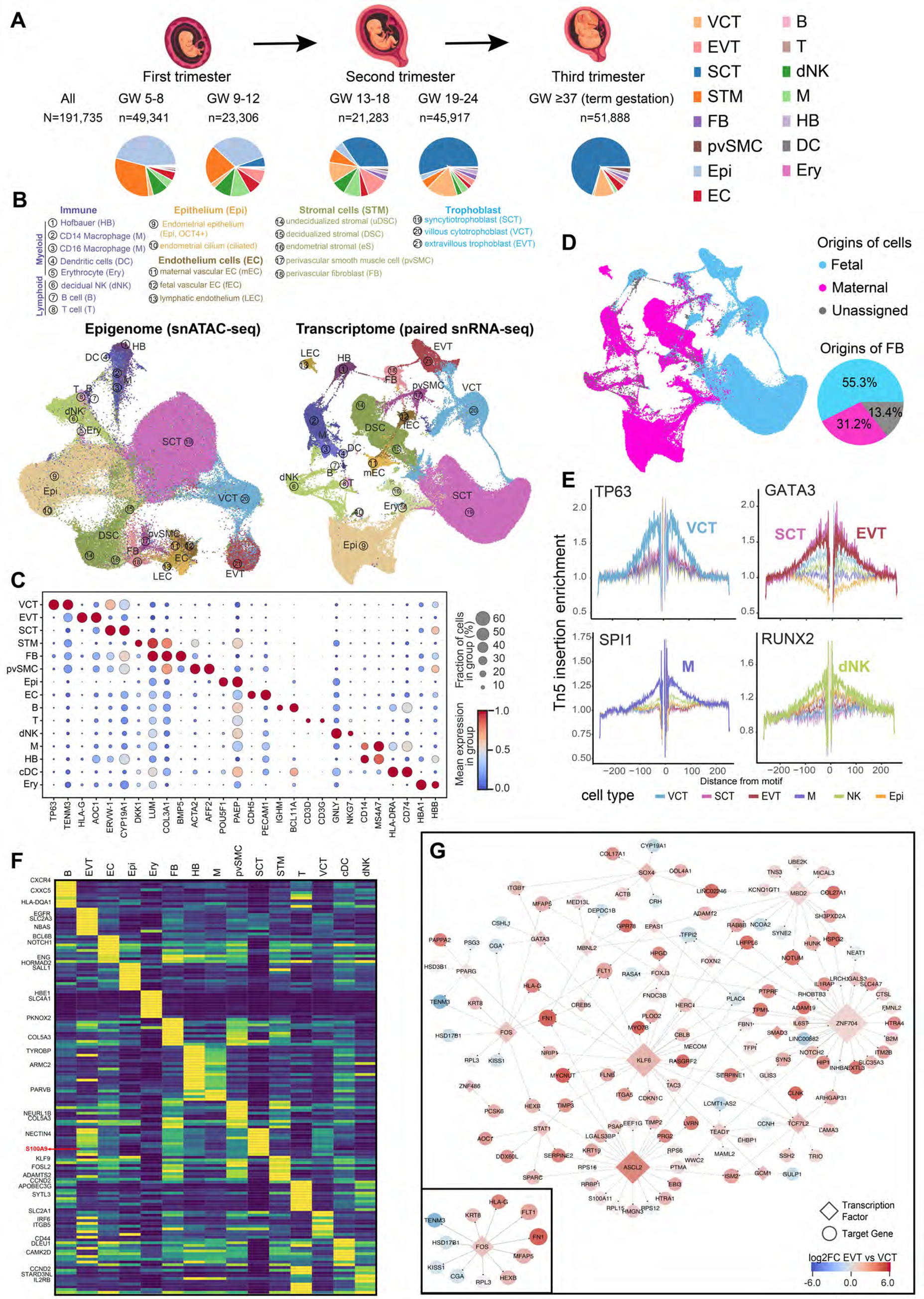
An overview of the single-cell multiomics data of the developing maternal-fetal interface. (A) Summary statistics of the samples, gestational weeks, the number of sequenced cells and the distribution of cell types across gestation. *N,* the total number of sequenced cells (post QC) in this study. *n*, the number of sequenced cells (post QC) in each gestational week range. **(B)** UMAP views of the clustering structure of 191,735 cells in the paired epigenome and transcriptome spaces. Each cell cluster was annotated by type. **(C)** Cell type marker genes displayed highly specific expression patterns in their corresponding cell types. **(D)** Identification of maternal or fetal origins of the sequenced cells. The UMAP was based on single cell transcriptomic data. Some cell types, such as mesenchymal cells, had mixed fetal and maternal origins. **(E)** Footprinting analysis of single cell ATAC-seq data identified physical binding sites of the major transcription factors in each cell type. Transcription factor binding sites were indicated by depletion of chromatin accessibility signals surrounding the binding motifs. **(F)** Genes with cell type specific enhancers in their *cis*-regulatory regions. S100A9 was highlight for its cell-type-specific enhancer in the SCT cell type. **(G)** Reconstruction of the gene regulatory network underlying the differentiation of villous to extra-villous cytotrophoblasts (VCTs->EVTs). Red and blue nodes are those with expression activation and repression, respectively, during the differentiation process. The inset shows the FOS-mediated regulon encompassing known cell markers or genes specifically expressed in EVT.

## Results

### Overview: Single Nucleus Multiomics Map of the Human Maternal-Fetal Interface

Our study design targeted cells at the human maternal-fetal interface from known normal (term gestation) or presumed normal (early/mid gestation) pregnancies. We included samples that did not have pregnancy complications and the included samples had no known fetal chromosomal or anatomic abnormalities (early-mid gestation). We collected fresh samples, that were rapidly snap frozen, from across gestation and used the decidua that included embedded trophoblasts (early gestation) or the basal plate (mid/late gestation) for our single cell multiomics profiling (**Fig.1A**). For early gestation samples, we confirmed collection of the decidua basalis vs. the decidua capsularis or parietalis by histology (**Fig.S1**) and immunolocalization of CTBs (**Fig.S2**). Because numerous placental chorionic villi (floating and anchoring) were associated with the decidua and basal plate, our sampling also captured their component cells. We performed 10X single-cell multiomics profiling, and generated post-QC, high-confidence, single-nucleus (sn) ATAC-seq and snRNA-seq data for 210,191 and 221,380 cells, respectively. Matching cell barcodes, we paired the data for 191,735 cells for downstream analyses, where each cell was simultaneously profiled by scATAC-seq and scRNA-seq and passed our rigorous 10X multiome QC analysis (**Methods and Materials, Table S2**). The number of sequenced cells across GW intervals is shown in **Fig.1A**. On average, 8,336 cells (post-QC) were multiomically profiled in each sample. Our subsequent analyses identified cell clusters in the epigenome (snATAC-seq) and transcriptome (snRNA-seq) spaces (**Fig.1B**). We annotated the clusters based on a panel of curated marker genes (**Fig.1C, Fig. S3, and Table S3**), which as expected showed strong cell-type specificities confirming the high quality of our data. The cell type composition of the samples is shown in **Fig.1A**. Due to inevitable sample variability as the cellular composition of the maternal-fetal interface evolves during gestation, we did not compare cell type composition across samples. Unless otherwise noted, we focused on the molecular characteristics of single cells.

To distinguish maternal vs. fetal cells, we implemented the Souporcell program(Heaton et al., 2020), which resolved the maternal or fetal origin of >95% of the sequenced cells (182,773/191,735, **Fig.1D**, see Methods and Materials). The accuracy was independently validated by Y-chromosome markers (**Fig.S4**, see Methods and Materials). We observed that several cell types, such as fibroblasts (FB), contained both maternal and fetal cells (**Fig.1D)**. This contrasted with vascular endothelial cells forming separate clusters for maternal and fetal cells (**Fig.1B**), suggesting distinct molecular characteristics. The paired epigenome and transcriptome data for each cell enabled a cell type comparison of the two spaces. Overall, cell type annotations in the epigenome and transcriptome spaces were concordant (**Fig.S5**), suggesting an epigenomic determination and subsequent transcriptomic manifestation of cell types at the maternal-fetal interface. Within each cell type, subtypes often exhibited strong separation in the transcriptome space; however, their boundaries were less clear in the epigenome space. For example, maternal lymphatic endothelial cells showed clear separation from maternal vascular endothelial cells in the transcriptome space (**Fig.1B**); but the two were not strongly differentiated in the epigenome space. The same was also true for fetal (Hofbauer cells) and maternal macrophages (**Fig.1B**). This observation suggested that at the maternal-fetal interface, the open-chromatin epigenome largely determines cell types at a high level, which, together with many other transcriptional regulatory mechanisms, ultimately defines cell subtypes at the transcriptomic level.

To investigate the molecular drivers of cell identity, we implemented chromVar(Schep et al., 2017) to infer transcription factors (TFs) that were enriched in each cell type. DNA binding motifs of functionally important TFs in a given cell type were expected to be significantly enriched in ATAC-seq peaks, suggesting their increased transcriptional activities. For example, the ATAC-seq peaks of VCTs were strongly enriched for TP63 an TFAP2B DNA motifs (**Fig.S6**); EVTs were enriched for TEAD1, GATA3 and GCM1 motifs (**Fig.S6**). snATAC-seq footprinting analysis(Stuart et al., 2021) of each cell type identified physical binding sites (∼20 base pairs in length) of TFs within pseudobulk ATAC-seq peaks, including TP63 (VCT), GATA3 (SCT and EVT), SPI1 (macrophages) and RUNX2 (natural killer [NK] cells) (**Fig.1E**).

Next, we mapped open chromatin ATAC-seq peaks onto *cis*-acting enhancer regions(Andersson et al., 2014) across the genome, which revealed cell type-specific active enhancer maps of cells at the maternal-fetal interface. Top 10 *cis*-acting enhancers specific to each cell type is shown in **Fig.1F (Table S4).** Of note, we identified a syncytiotrophoblast (SCT)- specific enhancer for *S100A9* (**Fig.1F**, red arrow)*, a* regulator of immune responses(Ryckman et al., 2003), whose expression was elevated in SCTs (**Fig.S7**). Dysregulation of S100A9 was recently associated with preeclampsia(Ozeki et al., 2022), highlighting the potential translational value of this enhancer map. When examining the EVT marker, *HLA-G*, which may play a role in maternal tolerance of the hemiallogeneic fetus(Hunt et al., 2005), we observed two significant ATAC-peaks specific to EVTs in addition to a promoter peak (**Fig.S8**). One ATAC-seq peak ∼10kb upstream of the *HLA-G* promoter corresponded to a recently reported active enhancer(Ferreira et al., 2016). A further upstream peak likely marks another novel distal enhancer (**Fig.S8**). The identification of cell type-specific *trans*-acting TF drivers and *cis*-acting enhancers revealed an extensive rewiring of the gene regulatory networks (GRN) that establish cell identities and their unique functions at the maternal-fetal interface.

To capture this network, we implemented CellOracle(Kamimoto et al., 2023) to integrate our single cell epigenomes and transcriptomes. For proof-of-principle, we reconstructed the GRN underlying the differentiation of VCTs to EVTs (**Fig.1G**), which physically and functionally anchor the placenta to the uterus(Red-Horse et al., 2004). Comparison of the VCT vs. the EVT transcriptomes identified 93 TFs that were significantly differentially expressed between the two cell types (FDR≤0.01). We identified the regulons of the 20 most highly dichotomous (minimal differential expression score greater than 37). Based on their cell-type-specific (correlated or anti-correlated) expression patterns and TF binding motif(s) in ATAC-seq peaks (± 250 bps of gene promoters), they mediated 167 high-confidence regulatory interactions (strength coefficient greater than 0.1) with 116 target genes (TGs). The GRN (**Fig.1G**) revealed several hub TFs mediating the most regulatory interactions, including ASCL2 and KLF6, known to regulate EVT differentiation(Miranda et al., 2022; Varberg et al., 2021), as well as ZNF704 and MBD2 (the NURD complex subunit) whose functions are not yet understood in this context. Notably, several genes whose expression were highly enriched in EVT were co-targeted by FOS, including the marker gene *HLA-G* (the best known EVT marker), KRT8 (**Fig.S9A**) and FN1 (fibronectin, **Fig.S9B**). In general, differentiation of VCTs to EVTs was associated with upregulations of TFs and their targets (red nodes, **Fig.1G**), but a few genes were significantly downregulated (blue nodes, **Fig.1G)**. Many of the latter were SCT markers: *CGA*, *PLAC4*, *CCNH*, and *TFPI2* (**Fig.S9C- G**). Thus, rewiring the transcriptional regulatory network to establish EVT identity requires simultaneously activating EVT-specific nodes *and* repressing genes expressed in other trophoblast lineages (*e.g.,* SCTs). This analysis, which revealed known and novel regulators of VCT differentiation into EVTs, shows the value of this resource for mapping rewired gene regulatory networks driving cell development at the maternal-fetal interface in normal pregnancy and aberrations thereof.

### Spatial Single-Cell Proteomics Profiling at the Maternal-Fetal Interface

Next, we localized the observed cell types from our single cell profiling (**Fig.1B**) at the maternal-fetal interface. We performed spatial single-cell proteomics profiling leveraging the CODEX technology (PhenoCycler, Akoya Biosciences), a multiplexed microfluidic immunofluorescence imaging system(Goltsev et al., 2018; Schurch et al., 2020). CODEX uses customizable panels of up to 50 antibodies conjugated to oligonucleotide barcode sequences in a single tissue staining reaction and achieves subcellular resolution. Guided by our single cell data, we examined the validated CODEX antibodies initially designed for the Human Tumor Atlas Network project(Rozenblatt-Rosen et al., 2020), and confirmed 9 antibodies specific to various cell type(s) at the maternal-fetal interface (**Table S5** and **Fig.2A**), including <u>anti-Ki67</u> marking proliferating cells (**Fig.S10**) and the other 8 shown in **Fig.2A-H:** <u>anti-CD3</u> (T cells); a<u>nti- E-cadherin</u> (CDH1, endometrial epithelium [strong expression] and VCT [moderate expression]); <u>anti-PDPN</u> (fibroblasts); <u>anti-VIM</u> (mesenchymal cells); <u>anti-CD31</u> (*PECAM1*; pan-endothelial); <u>anti-CK18</u> (pan-trophoblast with EVT having the strongest signal); <u>anti-CD206</u> (*MRC1*, maternal macrophages and fetal Hofbauer cells), and <u>anti-PDE3A</u> (arterial endothelium(Schupp et al., 2021) and perivascular smooth muscle cells(Omori and Kotera, 2007)). We particularly note PDE3A, whose expression in smooth muscle cells is known, and its specific expression in arterial endothelium was recently reported(Schupp et al., 2021). We further confirmed its specific expression in arterial endothelial cells using the Lung Endothelium Cell Atlas Data(Schupp et al., 2021) (**Fig.S11A**), our single-cell data in this study (described below) and our own immunolocalization experiments of uterine SAs (**Fig.S11B**). Because all our targeted cell types of interest were abundant at mid-gestation (**Fig.1A and 2A-H**), we performed CODEX profiling on two basal plate samples (GW 15.2 and 22.1, **Fig.2I** and **Fig.S12,** respectively), which yielded highly concordant results. Subsequent analyses focused on the GW15.2 sample that was richer in the cell types of interest. Data from the GW22.1 sample served as a replicate (**Fig.S12**).

**Fig. 2.**
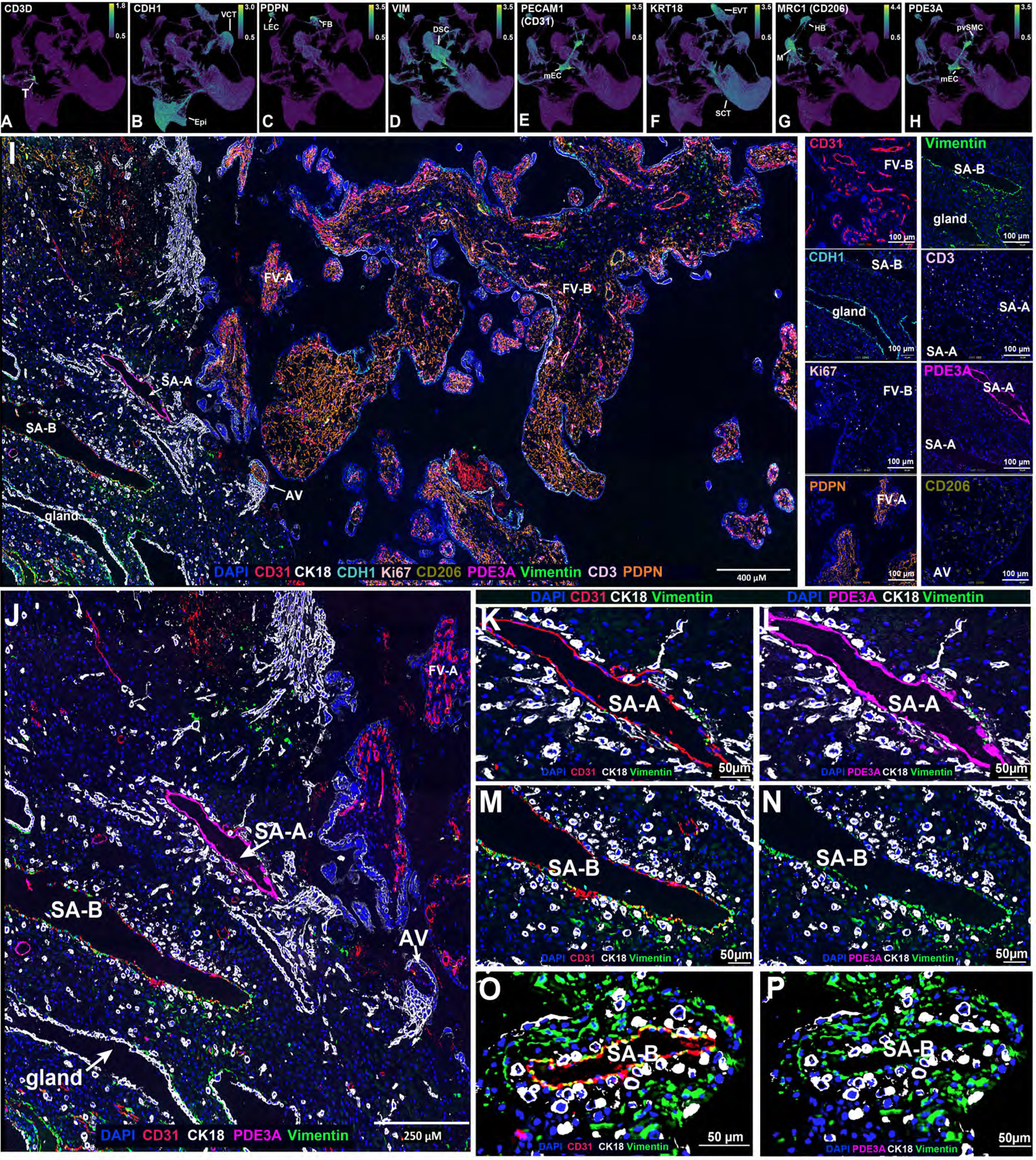
Single cell spatial proteomics profiling at the maternal-fetal interface using the CODEX technology. (A-H) The selected marker genes (and their specificity): CD3D (T cells), CDH1 (endometrial epithelium and trophoblasts), PDPN (lymphatic endothelia and mesenchymal stromal cells), VIM (stromal cell, and a subset of vascular and lymphatic endothelial cells), PECAM1 (CD31, endothelial cells), KRT18 (CK18, cytotrophoblasts, particularly EVTs), MRC1 (CD206, macrophages including fetal Hofbauer cells), and PDE3A (arterial endothelia and perivascular smooth muscle cells). **(I).** An overview of the CODEX multiplexed imaging data of 9 marker genes at the maternal-fetal interface (GW15.2). SA, spiral artery; AV, anchoring villi; FL, floating villi. Right panels display regional information: by viewing individual color channels, cells expressing a specific protein were readily observed, and cell types of interest were immunolocalized. **(J-P)** A high power view of two decidual SAs (SA-A and SA-B) revealed distinct molecular characteristics of their endothelial linings. **(O, P)** Analysis of an SA from a second basal plate sample (GW22.1) identified a vessel with the same endothelial molecular profile as SA-B (VIM^+^PDE3A^-^).

CODEX localized the major cell types identified in the single cell experiments (**Fig.2I; left panel**). Selective viewing of individual color channels revealed the distribution of specific cell types (**Fig.2I; right panels**). For example, in villous cores, we observed endothelial (CD31), Hofbauer (CD206) and mesenchymal (PDPN) cells. VCTs were marked by E-cadherin (CDH1, **Fig.2I** and **Fig.S13A, B**). Both VCTs and SCTs reacted with anti-CK18 (**Fig.2I, J**). However, the strongest CK18 staining was observed in EVTs, a result at the protein level that was highly consistent with our RNA data (**Fig.2F**). Uterine glandular epithelial cells also strongly reacted with anti-CK18 and anti-E-cadherin (**Fig.2I, J**). However, CK18^bright^ EVTs were easily identified by their association with anchoring villi, their interstitial distribution within the uterine wall and by their concentration around maternal blood vessels (**Fig.2I, J**). Combining data from multiple color channels enabled an investigation of communications among different cell types at the maternal-fetal interface. For example, within the uterine wall, T cells frequently surrounded EVTs (**Fig.S14**). We were particularly interested in EVT remodeling of SAs, which is one of the most intricate and critical steps in normal pregnancy and error prone in pregnancy complications(Labarrere et al., 2017; Staff et al., 2022). During this process, EVTs invade the vessel wall, intercalating within the muscular walls and displacing/replacing the resident endothelial cells. The net effect is conversion of these vessels to low resistance, high-capacity conduits. We investigated our CODEX data to study the molecular dynamics associated with SA remodeling by EVTs, followed by immunolocalization for verification and by single cell remapping for functional characterization.

Our CODEX profiling captured two intact SAs, which displayed highly contrasting patterns in their endothelial composition (SA-A and SA-B, **Fig.2I** and **J**). To investigate the communication between EVTs and endothelial cells, we filtered CODEX color channels associated with endothelial cells, including CD31 (marking endothelium), PDE3A (marking arterial endothelium), VIM (marking mesenchymal cells) and CK18 (marking EVTs in the decidua). Note that the two vessels had dense EVT clusters, suggesting active SA remodeling. Both SA-A and SA-B had CD31+ endothelial cells lining the vessel walls. A high-powered view revealed distinct molecular characteristics of the two SAs (**Fig.2J**). SA-A endothelial cells were negative for VIM but enriched for the arterial endothelial marker PDE3A (VIM^-^PDE3A^+^, **Fig.2K** and **L**). However, SA-B endothelial cells had the opposite expression pattern: positive for VIM and largely negative for PDE3A (VIM^+^PDE3A^-^ **Fig.2M** and **N)**. Individual PDE3A positive endothelial cells were also observed surrounding SA-B (**Fig.S13C**), suggesting a reminiscent signal associated with dynamic change of cell states which led to a loss of arterial cell identity and ultimately to apoptosis (see analysis below) due to SA remodulation.

To confirm that our observations were not specific to one sample, we performed CODEX profiling on another decidual biopsy (GW22.1) and identified one vessel of the SA-B type (VIM^+^PDE3A^-^) (**Fig.2O** and **P**). To replicate our observation of the SA-A-type (VIM^-^PED3A^+^), we performed immunolocalization on an GW20 decidual sample, and captured an arterial vessel of the SA-A type (low VIM and high PDE3A expression, VIM^-^PDE3A^+^) surrounded by EVT clusters (**Fig.3A-C**). Overall, these additional data indicated that the SA-A and SA-B phenotypes are commonly associated with EVT-mediated SA remodeling at the maternal-fetal interface.

**Fig. 3.**
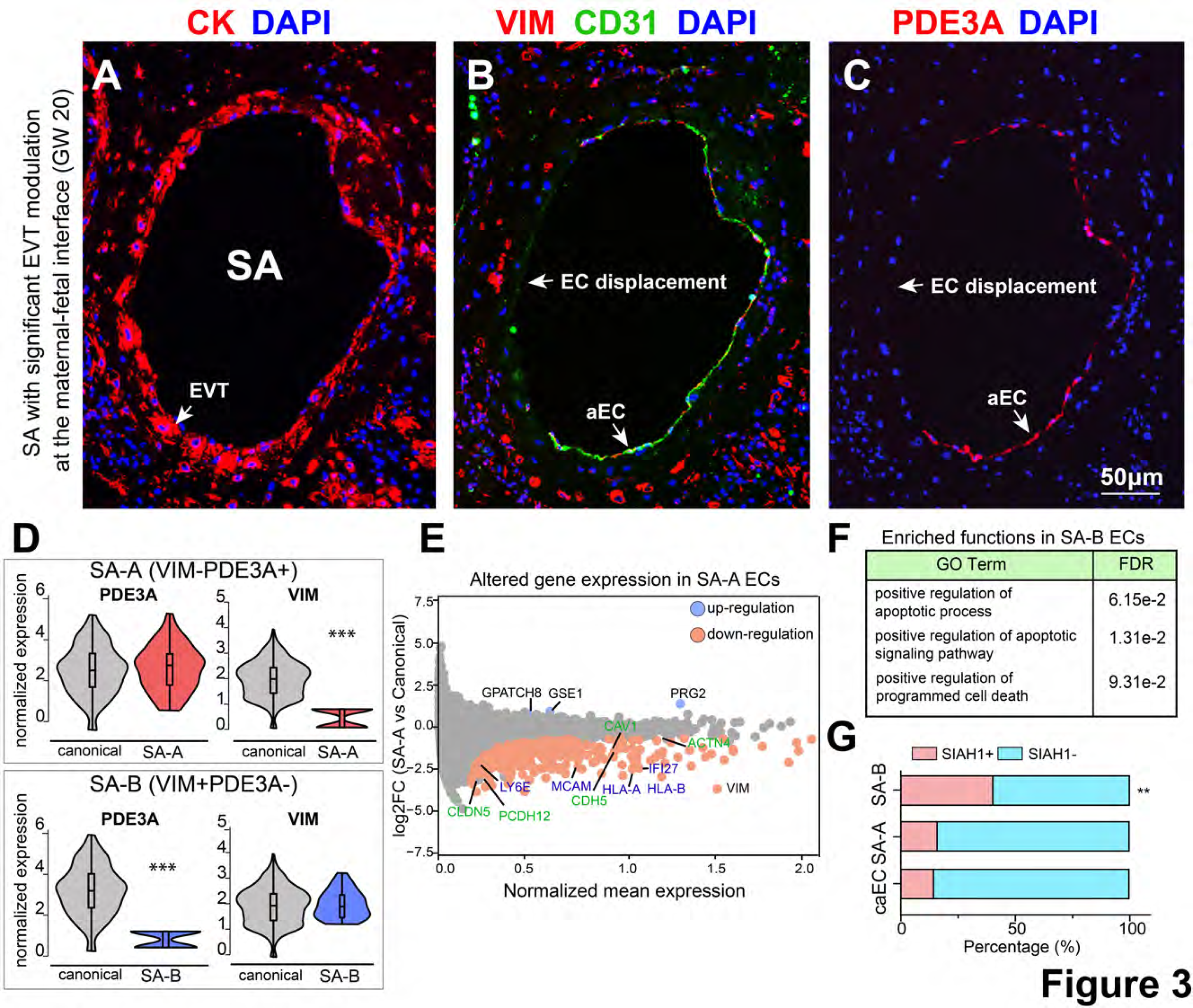
Distinctive molecular signatures of vascular endothelial states during SA remodeling. (A-C) Immunolocalization captured an SA undergoing EVT remodeling. Half of the arterial endothelial cells (aECs) had been displaced (EC displacement) by EVTs (marked by CK, pan-cytokeratin, **A**). The other half of the vessel was lined by aECs with the molecular profile of SA-A [(Fig.2) VIM^-^ (**B**), PDE3A^+^ (**C**)]. **(D)** Single cell data identified two groups of arterial endothelial cells: SA-A (VIM^-^PDE3A^+^) and SA-B (VIM^+^PDE3A^-^, Fig.2). All other arterial endothelial cells (aECs, X-axis) were used as canonical arterial endothelial cells for further functional comparisons. **(E)** Differentially expressed arterial endothelial genes in the SA-A vessel relative to canonical arterial endothelial cells. The downregulated genes in SA-A were enriched for cell junction (green) and immune (blue) genes. **(F)** Gene ontology enrichment of overrepresented genes in SA-B relative to canonical arterial endothelial cells. **(G)** The apoptosis regulator SIAH1 was highly enriched in SA-B cells relative to cells in SA-A or canonical arterial endothelial cells (caECs). P values were calculated by the Chi-square test. ** denotes a p < 0.01.

Because both VIM and PDE3A are present in canonical arterial endothelial cells (**Fig.S11**), losing either marker (SA-A- or SA-B-type, **Fig.2J-N**) suggests different endothelial states associated with EVT remodeling of SAs. We sought to identify these specialized cells leveraging our single cell data that included 3,514 maternal endothelial cells (**Fig.1B)**. We identified 497 maternal arterial endothelial cells highly enriched for known arterial endothelial cell markers (**Table S6**), where *PDE3A* clearly separated these cells from 3,017 maternal venous endothelial cells (**Fig.S15A-D**). Among 497 arterial endothelial cells, we identified 101 SA-A type cells (low VIM and high PDE3A [VIM^-^PDE3A^+^], **Fig.3D**) and 34 SA-B type cells (high VIM and low PDE3A [VIM^+^PDE3A^-^], **Fig.3D**). In comparison to all other arterial endothelial cells (the canonical endothelial cells, caECs), SA-A type cells displayed significant down-regulation of many genes implicated in immune responses, particularly those in antigen presentation (**Fig.3E, Table S7**). In addition, the down-regulated genes included those required for aspects of cell junction formation (**Fig.3E** and **Table S6**), including the tight junction (*CLDN5, ACTG1*), the gap junction (*CAV1*), the adherent junction (*CDH5, RAMP2*), as well as many related factors (*MCAM*, *CD99, PCDH12, ACTN4 and VIM*). These observations suggested immune desensitization and structural destabilization of the SA-A type cells (VIM^-^PDE3A^+^).

For the SA-B type cells (VIM^+^PDE3A^-^, **Fig.2K, L, O and P**), compared with canonical arterial endothelia, their loss of cell type was confirmed at the transcriptomic level by significant down- regulation of most arterial endothelial marker genes (**Fig.S15E, Table S6**). Across the transcriptome, we identified 61 genes upregulated in SA-B type cells than in canonical arterial endothelial cells (FDR<0.05) and the identified genes were uniquely enriched for regulating apoptosis (FDR=6.15e-2, **Fig.3F**). For example, the apoptosis regulator *SIAH1* was more likely to be expressed in these SA-B type cells than in canonical arterial endothelial cells (caECs, P=0.01, Chi-square test, **Fig.3G**), and the enrichment was absent for SA-A cells (P>0.1, Chi-square test, **Fig.3G**). Since these endothelial cells will eventually be displaced by EVT remodeling, this observation suggested that a pro-apoptotic state among the SA-B cells. Taken together, these data suggested that EVT remodeling of SAs results in at least two distinct endothelial states. The immune desensitization and structural destabilization of the SA-A type cells suggested cellular priming for their future displacement by EVTs, whereas the SA-B type cells were at a later apoptotic stage. These data were supported by our observation in **Fig.3A-C**: the partial loss of endothelial cells suggested priming of the remaining endothelial cells (SA-A) towards programmed cell death (SA-B).

### Reconstructing Cytotrophoblast Differentiation Trajectories

We performed UMAP clustering of 95,872 trophoblast cells. VCTs, SCTs, and EVTs – distinguished by expression of *TP63*, *HOPX* and *HLA-G*, respectively – formed unique clusters (**Fig.4A-C**). Each of these broadly defined cell types was comprised of multiple states **(Fig.4D**). To investigate the interrelationships, we used the Palantir method(Setty et al., 2019) to reconstruct their differentiation trajectories (**Fig.4E**). The results supported our current understanding of CTB differentiation: VCTs differentiate into SCTs or EVTs. However, advancing beyond previous single cell analyses, our profiling of more than 95,000 trophoblast cells across gestation identified many more cells at intermediate developmental stages, enabling high resolution to fine-map trophoblast differentiation trajectories and to uncover new cell (sub)types.

**Fig. 4.**
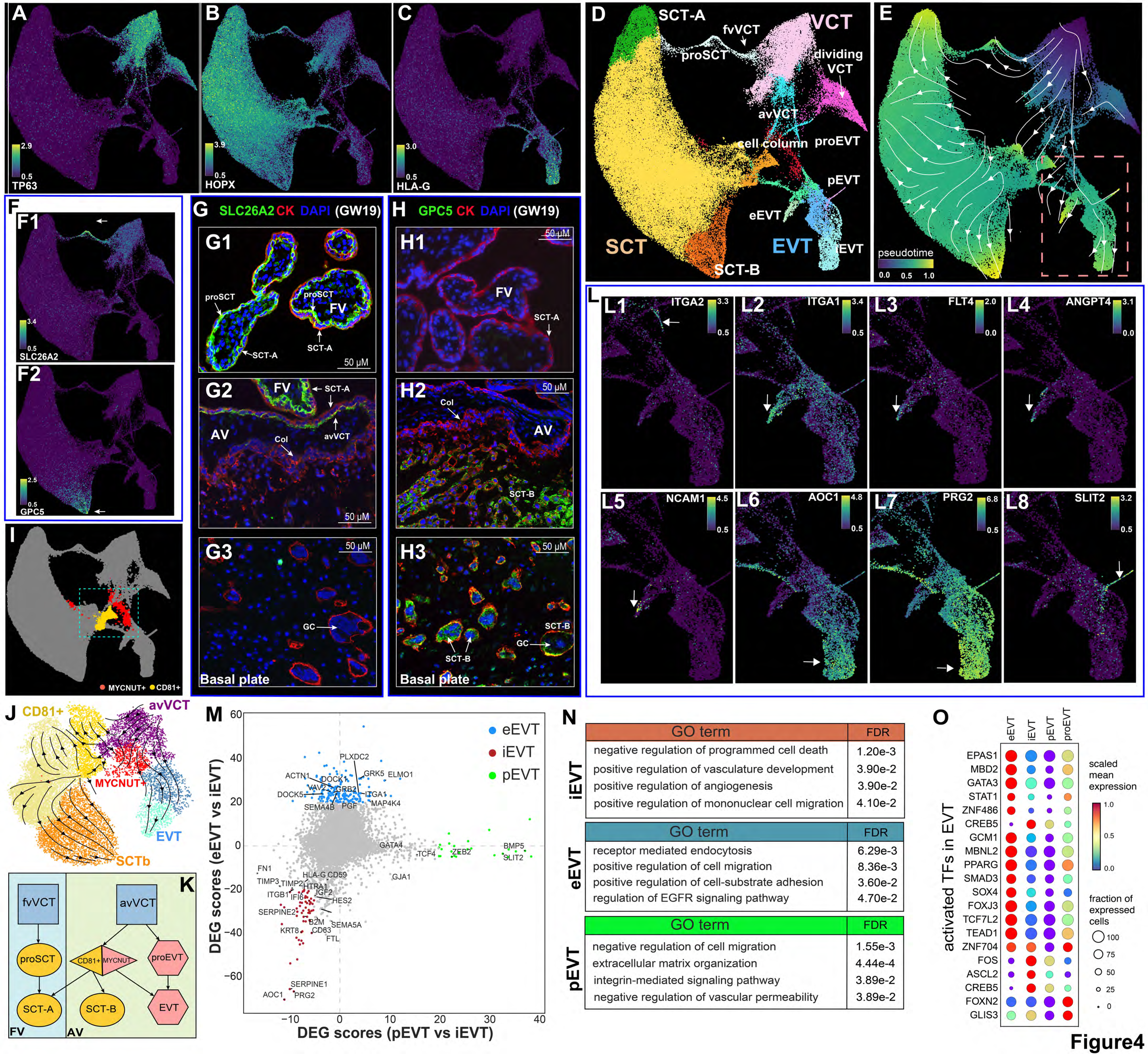
Molecular characterization of human trophoblast developmental trajectories. (A-C) UMAP projection for all trophoblast cells annotated with marker genes for VCTs (TP63), SCTs (HOPX), and EVTs (HLA-G). **(D)** UMAP projection of trophoblasts. Individual cell states are denoted by different colors. The major intermediate and terminal states along their developmental trajectories were labeled. (**E**) The reconstructed trophoblast differentiation trajectory revealed different developmental paths in floating and anchoring villi. Developmental pseudotime: purple (early stages) and yellow (late stages). (**F1)** Expression of SLC26A2 marked a VCT subtype (fvVCTs) that differentiated into a SCT subtype (SCT-A). (**F2**) GPC5 expression marks the other SCT subtype (SCT-B). **(G1-G3,** GW19.2) Immunolocalization confirmed the specificity of SLC26A2 as a marker for CK^+^ SCT-A (**G1**) in floating villi vs. anchoring villi (AV) (**G2**). CK^+^ column CTBs (**G2**), EVTs and trophoblast giant cells (GC) were also SLC26A2^-^ (**G3**). **(H1-H3,** GW19.2) CK^+^ trophoblast cells in floating villi were GPC5^-^. Immunostaining for this marker was negative in floating villi (**H1**) and was first detected among cytotrophoblasts (CTBs) leaving the cell column (**H2**). The strongest staining was in association with the multi-nucleated trophoblast giant cells (GC) in the basal plate (**H3**). (**I**) CD81 and MYCNUT expression stratified column CTBs into two intermediate cell states. (**J**) CD81^+^ cells differentiated into SCT-Bs, and MYCNUT^+^ cells differentiated into EVTs. **(K)** A model for trophoblast differentiation in floating villi (FV) or anchoring villi (AV). (**L1-L8**) Expression specificities of exemplar marker genes in EVTs and their subtypes. iEVT, interstitial EVT; eEVT, endovascular EVT; pEVT, perivascular EVT. **(M)** Differential gene expression analysis identified signature genes in the three EVT subtypes (color coded), and bkgd denotes the transcriptome background. DEG scores quantified differential gene expression among groups by taking into account the fraction of cells expressing a given gene and the fold changes among cell subtypes. **(N)** Enriched functional terms from gene ontology analysis for the signature gene sets of EVT subtypes. **(O)** Transcription factors (TFs) expression in the EVT subtypes as a function of VCT differentiation to EVTs. Shown are the top 20 TFs with the strongest activation.

Despite extreme cellular heterogeneity, we observed two terminal SCT states identified at the maternal-fetal interface (SCT-A and SCT-B, **Fig.4D** and **E**), each originating from a distinct VCT subtype (**Fig.4D**). Gene expression analysis identified *SLC26A2,* a sulfate transporter, as uniquely marking VCT and the transition from VCT to SCT-A (**Fig.4F1**). In contrast, *GPC5*, which encodes a cell surface heparan sulfate proteoglycan, uniquely marked the other SCT terminal state, SCT-B (**Fig.4F2**). We immunolocalized these SCT subtypes. We used SLC26A2 and cytokeratin (pan-trophoblast antigen) as makers for SCT-A at mid-gestation (GW=19.1; **Fig.4G**). In floating villi, anti-SLC26A2 stained VCTs. The signal was preferentially associated with the VCT cell surface adjacent to the syncytium, suggesting that these cells could be intermediate progenitors (proSCT) of SCT (SCT-A, **Fig.4G1**) as we also observed from single cell data (**Fig.4F1**). Clearly distinguishable SCTs at the surface of anchoring villi lacked or displayed low levels of SLC26A2 immunoreactivity (**Fig.4G2**), which was not detected in association with cell column cytotrophoblasts (**Fig.4G2**), or fused trophoblast giant cells (GCs) within the decidua (**Fig.4G3)**. Taken together, our data restricts the generation of SCT-A to a particular VCT subtype in floating villi (fvVCT, **Fig.4D**). Note that we observed that *SLC26A2* was co-expressed with *ERVFRD*-1 (*syncytin-2*), consistent with previous data suggesting that syncytin-2 rather than syncytin-1 promotes VCT fusion into multinucleated SCTs(Lokossou et al., 2014).

Next, we immunolocalized GPC5, which uniquely marked SCT-B (**Fig.4D** and **F2**). CK^+^ trophoblasts of floating villi as well as the syncytial surface of anchoring villi did not react with anti- GPC5 (**Fig.4H1**). In contrast, anchoring villi gave rise to CK^+^ trophoblast cells with increased anti- GPC5 immunoreactivity as they migrated from cell columns into the decidua (**Fig.4H2**). In deeper regions of the basal plate, multi-nucleated CK^+^ trophoblast giant cells also displayed strong GPC5 signals (**Fig.4H3**). The specificity of GPC5 expression by this SCT subtype (**Fig.4F2**) enabled identification of this novel subpopulation in cell columns and among CK^+^ cells migrating towards and eventually residing in the decidua. These data expanded our understanding of trophoblast giant cell (GCs) formation from fused EVTs to including also an important contribution from SCTs.

As the GPC5^+^ SCT-B cells migrate through anchoring villi, their developmental origin was traced back to a specific VCT subtype (avVCT), which also gave rise to EVTs (**Fig.4D** and **E**). As such, the differentiation from avVCT to SCT-B and to EVT delineated a distinct trajectory in anchoring villi vs. floating villi (from fvVCT to SCT-A, **Fig.4D** and **E**). Focusing on the intermediate trophoblast cell states in anchoring villi (the boxed area, **Fig.4I**), our transcriptomic analysis fine-mapped two subpopulations stratified by *CD81* and *MYCNUT* expression, differentiating into the SCT-B and EVT lineages, respectively (**Fig.4J**). Specifically, the *CD81*^+^ population was highly enriched for SCT marker genes and *MYCNUT*^+^ cells for EVT marker (**Fig.S16C-F**). Each population had a high probability of assuming the fate we ascribed (P=6.25e-141 and 1.35e-224, **Fig.4J** and **Fig.S16G-H**). This observation was also consistent with a previous study where CD81^+^ CTBs were identified as SCT progenitors, and their expression was specific to the proximal columns of anchoring villi(Shen et al., 2017). Notably, VCTs in anchoring villi also generated SCT- A (**Fig.4E**), which likely explains the origin of the syncytium that often forms the outer surface of anchoring villi. Taken together, these results revealed different developmental trajectories of VCTs in floating vs. anchoring villi (**Fig.4K**).

We were particularly interested in EVTs due to their significant roles in pregnancy outcomes. Expression of *ITGA2* marked EVT progenitors (proEVT) that were the direct descendants of VCTs in anchoring villi (**Fig.4D** and **L1**), an observation that was previously reported(Lee et al., 2018). As expected, mature EVTs expressed *HLA-G* (**Fig.4A**) and *ITGA1* (**Fig.4L2**). Trajectory analysis identified three terminal EVT states, corresponding to the interstitial (iEVT), endovascular (eEVT) and perivascular EVT (pEVT) subpopulations (**Fig.4D** and **E**). Specifically, we captured 1,014 cells that expressed well-known eEVT markers, including *FLT4*(**Fig.4L3**)(Zhou et al., 2002), *ANGPT4*(**Fig.4L4**)(Sato et al., 2003) and *NCAM1*(**Fig.4L5**)(Burrows et al., 1994). We captured 2,576 iEVTs. This subpopulation, which is distinguished by *AOC1* (**Fig.4L6**) and *PRG2* (**Fig.4L7**) expression(Haider et al., 2022), invades the decidua and reaches the inner third of the myometrium. Finally, we captured 221 pEVTs that specifically expressed genes regulating perivascular physiologies (described below), exemplified by *SLIT2* (**Fig.4L8**). Defining these subpopulations has enabled refining the expression patterns of the pan-EVT markers – notably, enriched expression of *HLA-G* in iEVTs (**Fig.4C** and **Fig.S17**) and of *ITGA1* in eEVTs (**Fig.4L2**).

We compared the transcriptomes among the three EVT subtypes using the iEVT transcriptome as a reference (see **Methods and Materials**). Differential expression analysis revealed molecular signatures of the EVT subtypes: 110, 170 and 125 significant genes in iEVTs, eEVTs, and pEVTs, respectively (**Fig.4M**, and **Table S8**). The iEVT signature was enriched for positive regulation of mononuclear cell migration (FDR=4.10e-2), positive regulation of vasculature development (FDR=3.90e-2) and regulation of angiogenesis (FDR=2.80e-2; **Fig.4N** and **Table S9**). These enrichments resulted from significant up-regulation of iEVT signature genes (*SERPINE1*, *FLT1*, *ANGPTL4* and *SEMA5A*), suggesting a role for interstitial EVTs in regulating uterine blood vessel development during pregnancy. iEVT signature genes were also enriched for negative regulation of programmed cell death (FDR=1.20e-3, **Fig.4N**), suggesting long-term survival of these cells across gestation.

eEVT signature genes were enriched for those regulating epithelial cell migration (FDR=1.16e-2, **Fig**.**4N**), cell-substrate adhesion (FDR=3.60e-2), and endothelial growth factor receptor signaling (FDR=4.70e-2, **Fig**.**4N**). These observations confirmed the known vascular functions of eEVTs. Querying the Mouse Genome Informatics database(Blake et al., 2021) for mutants of these eEVT signature genes revealed that they were enriched for phenotypes including: hemorrhage (FDR=1.31e-3), embryonic lethality (FDR=7.9e-3), embryonic growth retardation (FDR=2.00e-2), abnormal angiogenesis (FDR=2.02e-2) as well as numerous birth defects (open neural tube and ventricular septal defect, FDRs<0.01). These studies highlighted the physiological connections of eEVTs with abnormalities in placental and early fetal development.

Genes specific to pEVTs were enriched for negative regulation of cell migration (FDR=1.55e- 3, **Fig**.**4N**), integrin-mediated signaling (FDR=3.89e-2), and ECM organization (FDR=4.44e-4). Proteins in many of the associated GO terms were significantly associated with arterial stiffness based on genome-wide association analyses from the UK Biobank(Sudlow et al., 2015) (FDR=1.45e-17, odds ratio=5.42). These observations suggested specialized roles of pEVTs in modulating SA functions such as compliance.

We analyzed top TFs that were significantly regulated during VCT differentiation into EVTs (**Fig.1G, Methods and Materials**). Despite their overall increased expression in the EVT lineage, we observed their expression preference in specific EVT subtypes (EVT progenitors, eEVTs, iEVTs and pEVTs): their relative abundance displayed an overall enrichment in eEVTs, suggesting extensive regulatory dynamics to shape the eEVT transcriptome. In contrast, these TFs had much reduced expression in pEVTs, suggesting a more divergent regulatory network from other subtypes. This is reflected by their spread towards perivascular fibroblast cells from the EVT cluster in the transcriptome space (**Fig.S18**). Thus, in addition to the well-known eEVT mimicry of endothelial cells(Zhou et al., 1997), our results suggest that pEVTs bear a strong resemblance to the arterial pericyte layer.

### Resolving the Heterogeneity of Decidual Stromal Cells

Maternal decidual stromal cell (DSC) interactions with fetal trophoblasts are crucial for the initiation and maintenance of a normal pregnancy(Hess et al., 2007; Vinketova et al., 2016). However, given the extreme heterogeneity of DSCs, their functions are incompletely understood. Our single cell profiling captured 20,579 DSCs across gestation (**Fig.1B**). Subsequent UMAP projection of these cells revealed 5 distinct clusters (DSC0-DSC4, **Fig.5A**). We performed single cell transcriptome analysis to derive the full cell development trajectories and identified two main paths: <u>(Path A) DSC0→DSC1→DSC3</u>, and <u>(Path B) DSC0→DSC2→DSC4 (</u>**Fig.5B**). These trajectories were rooted at DSC0, which were most abundant during the first trimester (**Fig.5C**). In contrast, two of the mature progenies – DSC3 and DSC4 – were depleted at early gestation but became abundant at mid or late gestation (**Fig.5C**). To add further support, we examined key DSC marker genes. DSC0 cells had enriched expression of the undecidualized cell marker *ACTA2* (***SM α-ACTIN***, **Fig.5D1**) and low expression of the markers for the fully decidualized cells, i.e. *IGFBP1* (**Fig.5D2**) and *PRL* (*PROLACTIN*, **Fig 5D3**). These observations confirmed that DSC0 cells were undecidualized. At the termini, DSC3 and DSC4 had opposite gene signatures: low *ACTA2* expression (**Fig.5D1**) and enhanced *IGFBP1* (**Fig.5D2**) and *PRL* expression (**Fig.5D3**), consistent with their maturation and decidualized status.

**Fig. 5.**
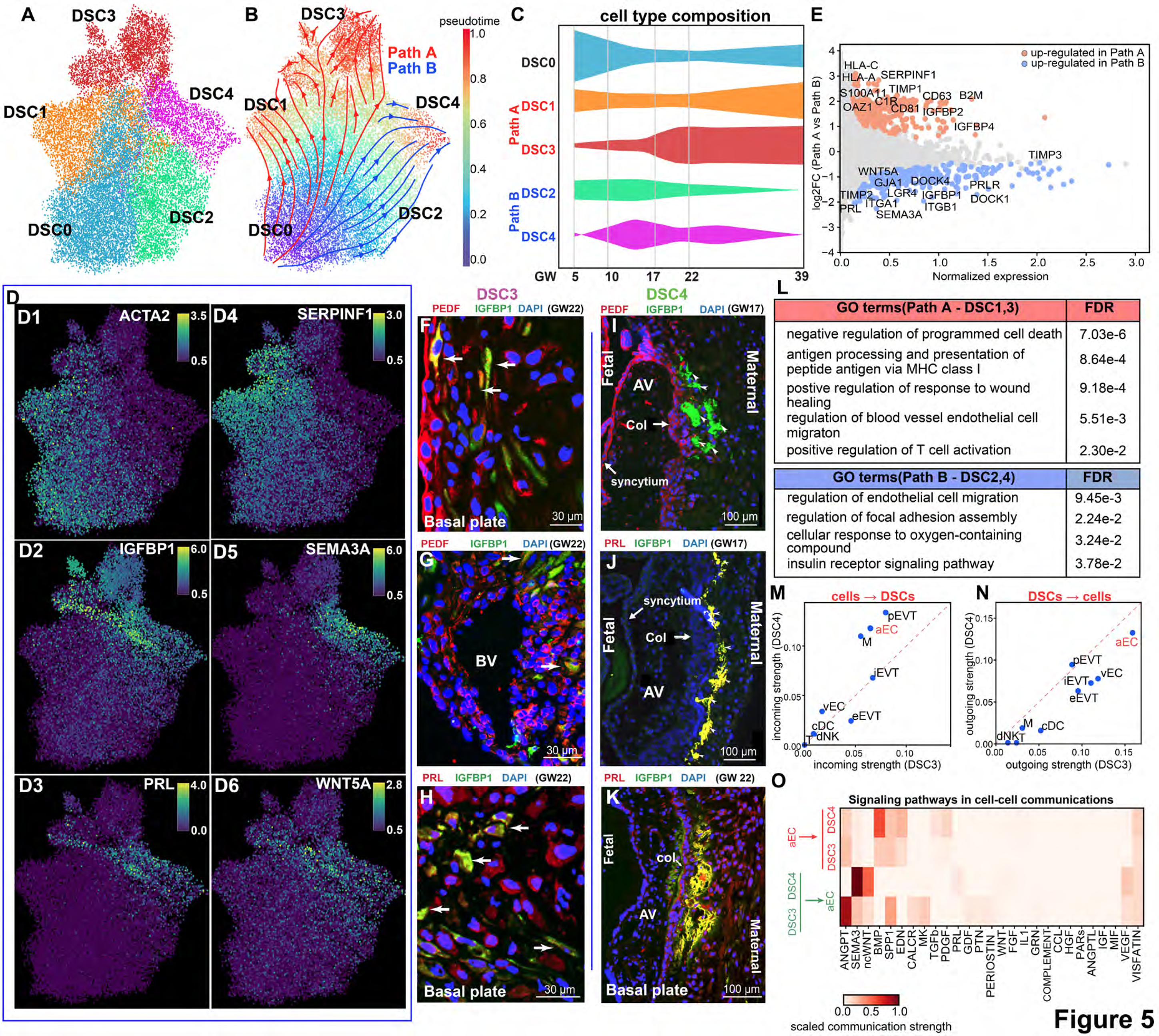
Resolving the heterogeneity of decidual stromal cells (DSCs). (A) Clustering analysis identified 6 subtypes of decidual stromal cells (DSC 0-5). **(B)** Two main trajectories were reconstructed underlying DSC development (early pregnancy to term): Path A: DSC0->DSC1->DSC3 and Path B: DSC0->DSC2->DSC4. **(C)** Percentage of cells in each DSC subtype across GWs. **(D)** The selected marker genes (and their specificity): ACTA2 (undecidualized cells, **D1**), IGFBP1 and PRL (decidualized cells, **D2** and **D3**), SERPINF1 (PATH A, **D4**), SEMA3A and WNT5A (PATH B, **D5** and **D6**). **(E)** Differential gene expression analysis identified SERPINF1 highly differentially expressed between the two developmental pathways (Path A and B). (**F-H,** GW22) DSC3 cells, co-stained with *SERPINF1* (PEDF) and IGFBP1, were located in the interstitial decidual regions **(F)** and sometimes along blood vessels (BV, **G**) in the basal plate. Their decidualized status was further confirmed by their co-expression IGFBP1 and PRL (**H**). (**I-K,** GW17 & GW22) Immunolocalization showed that DSC4 cells were confined to the regions immediately adjacent to the termini of anchoring villi. (**I**) DSC4 cells were PEDF^-^ and (**J-K**) IGFBP1^+^PRL^+^. AV, anchoring villi; Col, cell columns. (**L**) Enriched functional terms for the differentially expressed genes in Path A, Path B and Path C. (**M-N**). The strength of incoming (**M**) and outgoing (**N**) interactions between DSC3/4 and other cell types at the maternal fetal interfaces. Incoming interactions are other cell types (ligands) regulating DSC3/4 (receptors), and outgoing interactions are DSC3/4 (ligands) regulating other cell types (receptors). Above the diagonal line are cell types associated with stronger interaction strength with DSC4, and those below the diagonal line are associated with stronger interaction strength with DSC3. (**O**) The signaling pathways that contributed to arterial endothelium (aEC) regulation on DSC3 and DSC4 (the first two rows) and to DSC3/4 regulation to aEC (the last two rows).

We identified differentially expressed genes between Path A (DSC0→DSC1→DSC3) and Path B (DSC0→DSC2→DSC4, **Fig.5E**). *SERPINF1* (the PEDF protein, a potent antiangiogenic molecule(Dawson et al., 1999)) was specifically expressed in Path A vs. Path B (**Fig.5D4** and **E**). This result enabled immunolocalization of the terminally differentiated DSC3 (PEDF^+^IGFBP1^+^PRL^+^) and DSC4 (PEDF^-^IGFBP1^+^PRL^+^) cells at the maternal-fetal interface. First, we localized the undecidualized DSC0 cells in the decidua, where they displayed strong reactivity with anti-ACTA2 (**Fig.S20A).** We then sought to localize the fully decidualized DSC3 (PEDF^+^IGFBP1^+^PRL^+^) and DSC4 (PEDF^-^IGFBP1^+^PRL^+^) subpopulations. We observed that DSC3 cells were dispersed in the decidua (PEDF^+^IGFBP1^+^, **Fig.5F**), sometimes along maternal blood vessels (**Fig.5G**). As expected, these IGFBP1-positive DSC3 cells co-stained with PRL (**Fig.5H**), confirming their PEDF^+^IGFBP1^+^PRL^+^ phenotype. Notably, we also observed another group of PEDF^+^ cells that did not express IGFBP1 (PEDF^+^IGFBP1^-^, **Fig.5F and G**). Additional experiments confirmed that they could be CK^+^ cytotrophoblasts that expressed PEDF but not IGFBP1 **(Fig.S20B-D)**. Note that DSCs at an intermediate stage along Path A (DSC1, **Fig.5B**) could also express PEDF, and then IGFBP1 and PRL when reaching maturation (**Fig.5D2-D4**).

Unexpectedly, the DSC4 subpopulation (PEDF^-^IGFBP1^+^PRL^+^), which localized to the decidual surface, was restricted to areas that were adjacent to anchoring villi (**Fig.5I-K** and **Fig.S21** for independent replications). Consistent with our single-cell data (**Fig. 5D4** and **E**), these DSC4 cells (PEDF^-^IGFBP1^+^PRL^+^) were negative for PEDF (**Fig.5I**) but positive for IGFBP1 and PRL (**Fig.5J** and **K**). Notably, on the fetal side, syncytial cells associated with anchoring villi moderately expressed PEDF (**Fig.5I**) and PRL (**Fig.5K**), which also stained for CK (**Fig.S20B, H**). On the maternal side, further experiments confirmed that the mononuclear cells that only stained for PRL (**Fig.5J, H** and **K**) could be CK positive EVTs (**Fig.S20E-H**). Taken together, these data validated our single cell observations and confirmed the DSC developmental trajectories where Path A and Path B give rise to divergent DSC subpopulations with distinct molecular profiles and niches.

We sought to functionally characterize the two DSC developmental pathways (Path A and B, **Fig.5E**). The upregulated genes in Path A relative to Path B (**Table S10)** were enriched for negative regulation of programmed cell death (FDR=7.03e-6, **Fig.5L**), indicating the long-term presence of the DSC3 cells. Given the anti-angiogenic nature of PEDF (*SERPINF1*), the marker protein for Path A, Path A cells were also enriched for regulating blood vessel endothelial migration (FDR=5.51e-3, **Fig.5L**), consistent with their localization surrounding blood vessels (**Fig.5G**). In addition, these cells were also enriched for wound healing (FDR=9.18e-4, **Fig.5L**), antigen presentation (FDR=8.64e-4, **Fig.5L**) and T cell activation (FDR=2.30e-2, **Fig.5L**). We concluded that DSC3 cells (PDEF^+^IGFBP1^+^PRL^+^) might regulate anti-angiogenic processes while having unique immune functions.

Close examination of the Path A terminus (DSC3) revealed a bifurcation into two sub-clusters (DSC3.1 and DSC3.2, **Fig.S22**). The two subtypes expressed *IGFBP1*, confirming their decidualized status. DSC3.1 cells were absent after GW22 while DSC3.2 cells were present at all the timepoints studied. The transient DSC3.1 subtype lost the expression of angiogenic inhibitors (e.g., SERPINF1, TIMP1, **Fig.S22**), complement proteins (*e.g.,* C1R, C1S, C3, **Fig.S22**), MHC class I molecules (HLA/B/C, **Fig.S22**), and the apoptotic repressors (*e.g.,* PPIA, UBB, APOE, **Fig.S22)**. These observations suggested degeneration and pruning of DSC3 cells after GW22.

DSC4 cells were enriched for genes involved regulation of insulin receptor signaling (FDR=3.78e-2, **Fig.5L**), suggesting they could react to metabolic signals. These cells had increased expression of genes in regulating endothelial cell migration and focal adhesion assembly (FDR=2.24e-2, **Fig.5L**). Because the DSC4 formation of multilayered structures juxtaposed to anchoring villi (**Fig.5I-K**), it is likely they limit EVT invasion. This is supported by their specific expression of the chemorepellent *SEMA3A*(Kashiwagi et al., 2005)(**Fig.5D5**) as well as of *WNT5A* which regulates cell invasiveness and motility(Weeraratna et al., 2002) (**Fig.5D6**). It is also important to highlight that the strongly expressed *IGFBP1* in DSC4 (**Fig.5I-K**) is known to inhibit EVT invasion by its interaction with the cytotrophoblast ɑ5β1 integrin(Irwin and Giudice, 1998). We further examined a large collection of single cell data from non-pregnant endometrial samples at both proliferative and secretory phases(Garcia-Alonso et al., 2021), and observed that DSCs in the nonpregnant samples were negative for the DSC4-specifiic gene *SEMA3A*. This observation suggests that the DSC4 subtype is specific to human pregnancy stages and is absent in the nonpregnant endometrium.

For independent validation, we performed single-cell RNA-seq of decidual samples from an additional cohort of 13 individuals (GW39, where DSC3 and DSC4 were most enriched, **Fig. 5C**, Methods and Materials), generating transcriptomic profiles of 28,626 cells. Again, the DSC cells were defined by the expression of *SERPINF1* (PEDF), recapitulating cells in Path A and B trajectories (**Fig.S23**). As expected, DSCs negative for *SERPINF1* (PEDF) specifically expressed *SEMA3A* and *WNT5A* (**Fig.S23**), recapitulating the cell markers of DSC4 (described above). This replication experiment suggested that our conclusions were broadly applicable. Moreover, the consistency of the single-cell RNA-seq and single nuclei data suggested that our results were not platform specific.

We performed cell-cell communication analysis to characterize interactions of the decidualized DSC3/4 cells with other cell types at the maternal-fetal interface, including arterial (aECs) and venous endothelial cells (vECs), EVT subtypes (pEVT, iEVT and eEVT), and the major immune cell types (macrophages [M], classical dendritic cells [cDC], T and decidual NK [dNK] cells). We implemented the widely used CellChat(Jin et al., 2021), which scores the strength of interactions between two cell types based on the reciprocal expression of all possible ligand-receptor gene pairs. DSC3 and DSC4 had increased interaction strength with aECs and EVTs, regardless of signal directionality (**Fig.5M** and **N**). As to influences from neighboring cells, most immune cells had attenuated interactions with DSC3 and DSC4. However, macrophages had significant roles in regulating DSCs (**Fig.5M**) but lacked the ability to respond to these cells (**Fig.5N**). Macrophages, aECs and pEVTs had stronger regulatory interactions with DSC4 vs. DSC3 (**Fig.5M**). DSC4 cells are juxtaposed to anchoring villi (**Fig.5I-K**) in the vicinity of SAs. In this context, the elevated regulatory strength of aECs and pEVTs on DSC4 indicated these cells as possible regulators of the latter. In contrast, the influence of venous endothelial cells (vEC) was marginal (**Fig.5M**). Compared to DSC4, DSC3 displayed stronger interaction strength in terms of regulating aEC, vEC, iEVT and eEVT; the strongest effect was on aECs (**Fig.5N**). Given the anti-angiogenic role of DSC3 and their peri-vascular location (**Fig.5G**), these observations highlighted the interplay among DSC, macrophage, EVT and SA aECs.

To identify the underlying pathways, we analyzed the cell-cell communication strength from 27 curated signaling pathways between aECs and two terminal DSC types (DSC3/4) using CellChat. The strong interaction of aEC on DSC4 mainly relies on the BMP signals. Additionally, DSC3 and DSC4 cells regulated aECs through distinct signaling cascades: the anti-angiogenic DSC3 subtype signals through the angiopoietin-Tie pathway (ANGPT), whereas DSC4 utilizes the class-3 semaphorin (SEMA3) and non-canonical WNT (ncWNT) signaling pathways (**Fig.5O**). Taken together, analyses of intercellular communication demonstrated divergent functions of the two DSC subtypes.

### Integrating GWAS and Single Cell Data Identified Affected Cell Types in Major Pregnancy Complications

To demonstrate the translational value of our single cell compilation, we leveraged the data to guide our analysis of large-scale patient genomes with major pregnancy complications: preeclampsia, spontaneous preterm births, and sporadic miscarriages. These conditions likely have a significant genetic basis, but genome-wide association studies (GWAS) have implicated very few genomic loci. Thus, their molecular etiologies remain largely elusive.

More than 90% of disease-associated variants fall in noncoding regions, resulting in epigenetic alterations in active regulatory elements and, consequently, aberrant gene expression(Corradin and Scacheri, 2014; Schaub et al., 2012). Thus, it is possible to determine whether a cell type-specific epigenome is enriched for disease-associated genomic variants. The recently developed SCAVENGE framework integrates GWAS with snATAC-seq data to discover disease-associated cell types(Yu et al., 2022). The algorithm identifies single cell epigenomes that are enriched for common variants with increased GWAS risk, revealing the cell types most vulnerable to a given disease.

First, we applied this approach to preeclampsia, a common and potentially life-threatening pregnancy complication that manifests as a sudden rise in maternal blood pressure and the appearance of protein in the urine. Both signs are the result of maternal vascular damage. This syndrome has a genetic component with heritability as high as 60%; 35%, and 20% are potentially contributed by the maternal and fetal genomes, respectively(Rana et al., 2019). We analyzed the largest GWAS preeclampsia study to date(Steinthorsdottir et al., 2020), and only considered subjects with European ancestry. This study utilized 10.8 million genomic variants genotyped in 10,255 maternal cases and 10,255 matched female controls, together with 10.4 million variants in 7,259 affected fetal genomes (offspring of preeclamptic pregnancies) and 7,259 matched fetal controls. A standard GWAS analysis yielded only two significant SNPs from the maternal genomes and one from the fetal genomes. Their moderate effects sizes (OR<1.3) cannot explain preeclampsia heritability observed in population studies.

To identify cellular origins of preeclampsia population genetic risk, we implemented SCAVENGE to integrate this preeclampsia GWAS dataset with our snATAC-seq map. We paired maternal genomes with maternal cells, and fetal genomes with fetal cells. SCAVENGE computed the trait relevance score (TRS) for each cell (**Fig. 6A**) and identified, by permutation analysis, 6,221 maternal and 8,232 fetal cells as significantly associated with preeclampsia. The cells (maternal and fetal) accounted for 7.5% of all those identified (14,453/191,735). Next, we performed Fisher’s exact test to determine the enrichment of preeclampsia-associated cells in each cell type (subtype) relative to their identity as maternal or fetal. Among all fetal cell types, only EVTs (all subtypes) displayed a significant enrichment for preeclampsia-associated cells (FDR≤1e-4, **Fig.6B**); iEVTs manifested the strongest enrichment. In contrast, multiple maternal cell types were significantly associated with the genetic risk of preeclampsia. Among the DSC subtypes (**Fig.5A**), only anti-angiogenic DSC3 cells, the terminal state of Path A (**Fig.5B**), displayed significant enrichment in preeclampsia (**Fig.6C**). A subset of vascular components also had significant risk enrichments: arterial endothelium (but not venous endothelium), perivascular smooth muscle and perivascular fibroblasts (**Fig.6C**). Moreover, the enrichment of T cells (**Fig.6C**) was evidence of maternal immune involvement. Despite the proposed role of decidual macrophage and NK cells in preeclampsia(Faas et al., 2014; Hiby et al., 2004), population genetic risk was not enriched in these cell types. The association of a unique population of endometrial epithelial cells was of particular interest (**Fig.6C**). Many of these cells significantly expressed stem cell markers (OCT4 [*POU5F1*] and LGR5, **Fig.S24**), and their presence was limited to early gestation samples. This finding associated early gestation events with genetic etiologies of preeclampsia and suggested an endometrial contribution to the origin of preeclampsia(Conrad, 2020).

**Fig. 6.**
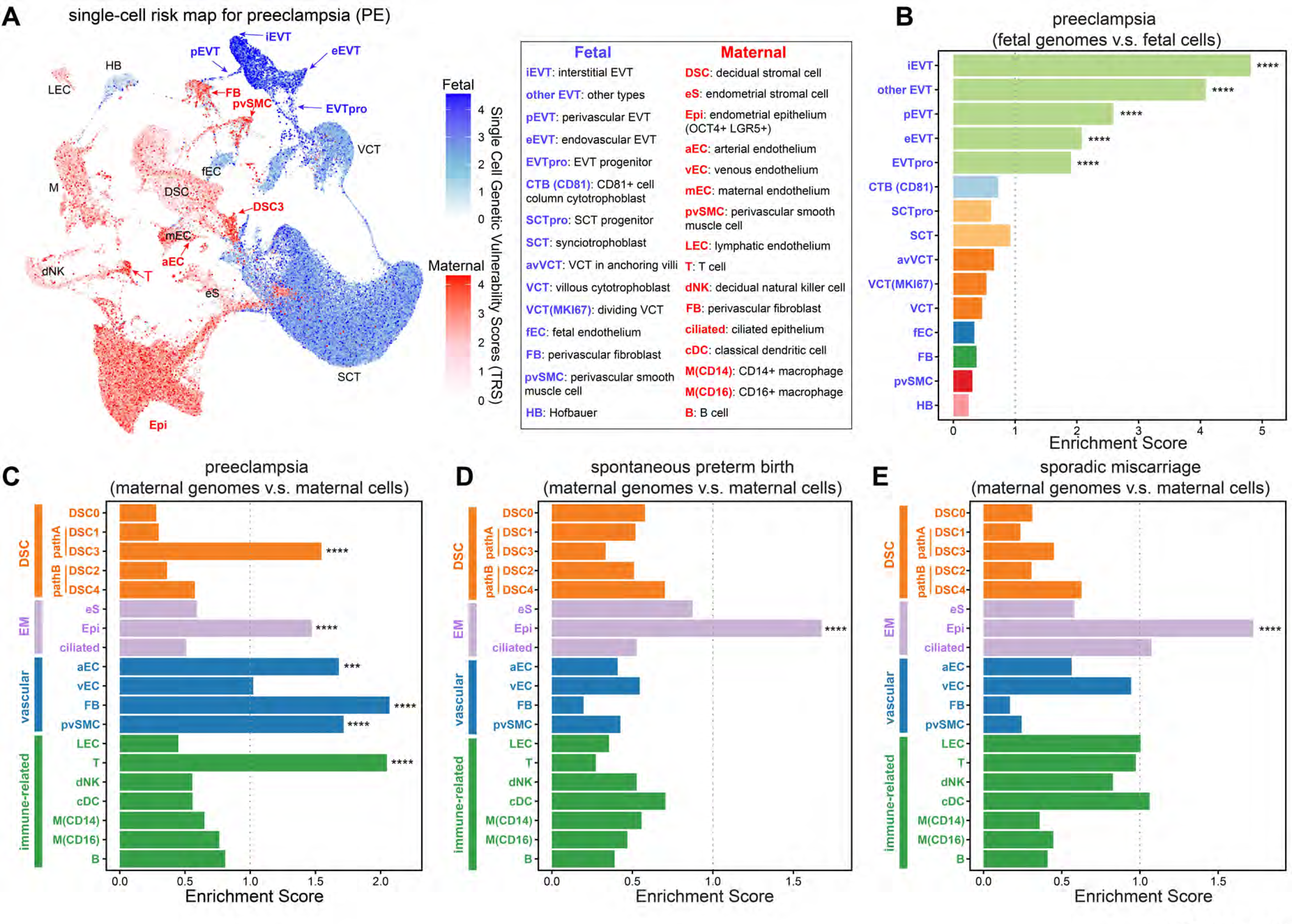
Constructing the single cell risk map for the major pregnancy complications. (A) Integrating single cell multiomics data with preeclampsia GWAS identified the cell types enriched for high-risk variants. Maternal and fetal GWAS were integrated with their respective cell types. **(B-C)** Cell (sub)types that were significantly associated with preeclampsia risk. Statistical significance was determined by Fisher’s exact tests. P values were adjusted by Benjamini- Hochberg correction. The percentage of significant cells in each cell type was compared with that of all cells (maternal or fetal), and the enrichment fold change was shown on X axis. **** indicated P<1e-4. Fetal (**B**) and maternal (**C**) GWAS preeclampsia associations with fetal and maternal cells, respectively. (**D-E**) The same analysis was performed for spontaneous preterm birth (**D**) and sporadic miscarriage (**E**). Subtypes less than 200 cells were excluded from the analyses.

The same analysis was performed for spontaneous preterm labor and spontaneous miscarriage. Because only maternal genomes were available for both conditions, we limited our analysis to maternal cell types. For spontaneous preterm labor, we analyzed GWAS data from 3,331 cases (GW<37) and 37,803 matched female controls(Zhang et al., 2017). For sporadic miscarriage, we analyzed GWAS data from 49,996 cases and 174,109 matched female controls(Laisk et al., 2020). In both cases, only the OCT4^+^LGR5^+^ endometrial epithelial cells displayed significant enrichment for genetic risk at a population level (**Fig.6 D** and **E**). This finding likely explains the clinical associations between miscarriage and spontaneous preterm labor (Oliver-Williams et al., 2015; Swingle et al., 2009). Given that this analysis also implicated these cells in preeclampsia, our results support the concept of “endometrium spectrum disorders”(Conrad et al., 2017), which suggests that many pregnancy complications are associated with a continuum of dysregulated endometrial functions. In this regard, we theorize that OCT4^+^LGR5^+^ epithelial cells play a particularly important deterministic role.

## Discussion

Here, we report the results of the most in-depth, single cell, multiomics analysis of the maternal-fetal interface. Integration across gestation, i.e., adding the additional dimension of development over time, resolved the extreme cellular heterogeneity inherent in two rapidly changing organs from two different individuals. This rich single cell genomic resource was the foundation on which we built our spatial single-cell proteomics profiling and functional analyses, which enabled the discovery of novel developmental trajectories and cell types. Furthermore, data integration with large-scale maternal and/or fetal genomes revealed the cell types at the maternal-fetal interface that are most vulnerable to the major pregnancy complications.

Our study leveraged the state-of-the-art CODEX platform to localize the major cell types at the maternal-fetal interface *in situ* (**Fig.2**). Compared with the recent spatial analysis of cytotrophoblasts using the 10X Visium platform(Arutyunyan et al., 2023) which has remained at a bulk tissue level (each spatial spot contains usually contains more than 10 cells), the CODEX system has achieved single-cell or even subcellular resolution (600nm or 250 nm). Compared with most spatial systems limited to gene expression (including 10X Visium), our CODEX profiling resolves protein abundance, enabling simultaneous localization of multiple cell types identified from our single cell sequencing. This unprecedented level of resolution revealed new information about the dynamic nature of SA endothelial cell responses to EVT remodeling (**Fig.2K-P)**. Specifically, endothelial cells lining SA-A type-vessels (**Fig.2J, M** and **N**) evidenced significant structural destabilization, which could prime the cells for subsequent apoptosis (SA-B). SA-A type endothelial cells also displayed highly significant losses of immune response genes (**Fig.3E**), particularly antigen-presenting molecules. This could be part of a strategy for suppressing maternal immunoreactivity against invading fetal EVTs. Based on these observations, we hypothesized that perturbations in the conversion of SA endothelium to either the SA-A or SA-B phenotype, could hinder EVT modulation as is observed in severe preeclampsia, fetal growth restriction and a subset of PTB cases. Finally, previous studies suggest that SA remodeling is accompanied by an arterial-to-venous fate transition of the resident endothelial cells(Pawlak et al., 2019; Zhang et al., 2008). This observation, based on the expression of individual genes, was consistent with our transcriptomic data: both SA-A and SA-B endothelial cells displayed down- regulation of arterial-type genes and up-regulation of venous-type genes (**Fig.S15**). However, their transcriptomes were still clustered with arterial endothelial cells rather than venous cells. Therefore, SA remolding likely precipitates alterations in their phenotype rather than fate.

Powered by our deep single cell multiomics data, we fine-mapped the developmental trajectories of trophoblast cells. The results added new knowledge to the paradigm that VCTs differentiate into SCTs or EVTs. Specifically, we show that VCTs in floating and anchoring villi have different trajectories. This finding supports a previous speculation that SCTs and EVTs originate from two distinct VCT subtypes(James et al., 2005). We verified that floating villi VCTs (fvVCTs) differentiate into SCTs. Unexpectedly, anchoring villi VCTs (avVCTs) gave rise to a novel SCT subtype (SCT-B, **Fig.4D**) that migrated through the cell columns and formed GCs that resided in the decidua. This finding contrasts with the assumption that GCs form via EVT fusion. Additionally, we showed that anchoring villi are crossroads where several different SCT and EVT developmental trajectories intersect **(Fig.4D** and **J**). We hypothesize that correct coordination of the differentiation pathways that unfold in this highly specialized placental region is a requisite for normal pregnancy. Conversely, deviations could be associated with pregnancy complications, such as preeclampsia, that negatively impact the formation EVTs. Our finding that genetic vulnerability of fetal cells to preeclampsia solely maps to EVTs supports this theory. Finally, it is important to note that our study only targeted the maternal-fetal interface. Many more trophoblast subtypes are likely to be identified in different placental and extraembryonic compartments, such as the recently identified subtype specific to the smooth chorion(Marsh et al., 2022).

Despite their heterogeneity, our single cell analysis revealed that DSCs converge onto common developmental trajectories. We showed that one subtype (DSC3) exclusively expressed the potent anti-angiogenic factor *SERPINF1* (PEDF) and is significantly associated with preeclampsia (**Fig.6C**). Immunolocalization showed that these cells resided in the decidua and perivascular regions of vessels that traversed this region. They degenerated after GW22, suggesting a gestational age-dependent mechanism for pruning anti-angiogenic DSCs prior to SA revascularization later in pregnancy. We hypothesize that, an overabundance of these cells or a defect in their pruning could be associated with preeclampsia, which is characterized by an anti-angiogenic state.

We discovered two new DSC subtypes. DSC4 cells expressed molecules that play key roles in inhibiting cell migration and invasiveness. Unexpectedly, they immunolocalized to the termini of anchoring villi where they formed multi-layered structures. Their phenotype and positioning strongly suggested that they counterbalance aggressive EVT penetration of the uterine wall. This is also reflected by the strong IGFBP1 expression in DSC4 (**Fig.5I-K**) which is known to inhibit EVT invasion (Irwin and Giudice, 1998).

Our rich single cell data provided us an opportunity to uncover vulnerable cell types to pregnancy complications. The identified cell types in pregnancy complications (preeclampsia, preterm births, and miscarriages) reflected genetic vulnerability at a population level (*i.e.,* common variants), which did not exclude contribution from rare mutations or from non-genetic factors. Overall, this study provides new insights into the coordinated development of the placenta and decidua across gestation in normal pregnancy and the cellular basis of pregnancy complications whose etiologies remain poorly understood.

## Acknowledgements

We acknowledge funding support from the Chan-Zuckerberg Initiative (the Silicon Valley Community Foundation) and NIH (NIHMS and NICHD)

## References

1. Andersson, R., Gebhard, C., Miguel-Escalada, I., Hoof, I., Bornholdt, J., Boyd, M., Chen, Y., Zhao, X., Schmidl, C., Suzuki, T., et al. (2014). An atlas of active enhancers across human cell types and tissues. Nature 507, 455–461.

2. Arutyunyan, A., Roberts, K., Troule, K., Wong, F.C.K., Sheridan, M.A., Kats, I., Garcia-Alonso, L., Velten, B., Hoo, R., Ruiz-Morales, E.R., et al. (2023). Spatial multiomics map of trophoblast development in early pregnancy. Nature 616, 143–151.

3. Blake, J.A., Baldarelli, R., Kadin, J.A., Richardson, J.E., Smith, C.L., Bult, C.J., and Mouse Genome Database, G. (2021). Mouse Genome Database (MGD): Knowledgebase for mouse-human comparative biology. Nucleic Acids Res 49, D981–D987.

4. Burrows, T.D., King, A., and Loke, Y.W. (1994). Expression of adhesion molecules by endovascular trophoblast and decidual endothelial cells: implications for vascular invasion during implantation. Placenta 15, 21–33.

5. Conrad, K.P. (2020). Evidence for Corpus Luteal and Endometrial Origins of Adverse Pregnancy Outcomes in Women Conceiving with or Without Assisted Reproduction. Obstet Gynecol Clin North Am 47, 163–181.

6. Conrad, K.P., Rabaglino, M.B., and Post Uiterweer, E.D. (2017). Emerging role for dysregulated decidualization in the genesis of preeclampsia. Placenta 60, 119–129.

7. Corradin, O., and Scacheri, P.C. (2014). Enhancer variants: evaluating functions in common disease. Genome Med 6, 85.

8. Dawson, D.W., Volpert, O.V., Gillis, P., Crawford, S.E., Xu, H., Benedict, W., and Bouck, N.P. (1999). Pigment epithelium-derived factor: a potent inhibitor of angiogenesis. Science 285, 245–248.

9. Faas, M.M., Spaans, F., and De Vos, P. (2014). Monocytes and macrophages in pregnancy and pre-eclampsia. Front Immunol 5, 298.

10. Ferreira, L.M., Meissner, T.B., Mikkelsen, T.S., Mallard, W., O’Donnell, C.W., Tilburgs, T., Gomes, H.A., Camahort, R., Sherwood, R.I., Gifford, D.K., et al. (2016). A distant trophoblast-specific enhancer controls HLA-G expression at the maternal-fetal interface. Proc Natl Acad Sci U S A 113, 5364–5369.

11. Garcia-Alonso, L., Handfield, L.F., Roberts, K., Nikolakopoulou, K., Fernando, R.C., Gardner, L., Woodhams, B., Arutyunyan, A., Polanski, K., Hoo, R., et al. (2021). Mapping the temporal and spatial dynamics of the human endometrium in vivo and in vitro. Nat Genet 53, 1698–1711.

12. Goltsev, Y., Samusik, N., Kennedy-Darling, J., Bhate, S., Hale, M., Vazquez, G., Black, S., and Nolan, G.P. (2018). Deep Profiling of Mouse Splenic Architecture with CODEX Multiplexed Imaging. Cell 174, 968–981 e915.

13. Greenbaum, S., Averbukh, I., Soon, E., Rizzuto, G., Baranski, A., Greenwald, N.F., Kagel, A., Bosse, M., Jaswa, E.G., Khair, Z., et al. (2023). A spatially resolved timeline of the human maternal-fetal interface. Nature 619, 595–605.

14. Haider, S., Lackner, A.I., Dietrich, B., Kunihs, V., Haslinger, P., Meinhardt, G., Maxian, T., Saleh, L., Fiala, C., Pollheimer, J., et al. (2022). Transforming growth factor-beta signaling governs the differentiation program of extravillous trophoblasts in the developing human placenta. Proc Natl Acad Sci U S A 119, e2120667119.

15. Heaton, H., Talman, A.M., Knights, A., Imaz, M., Gaffney, D.J., Durbin, R., Hemberg, M., and Lawniczak, M.K.N. (2020). Souporcell: robust clustering of single-cell RNA-seq data by genotype without reference genotypes. Nat Methods 17, 615–620.

16. Hess, A.P., Hamilton, A.E., Talbi, S., Dosiou, C., Nyegaard, M., Nayak, N., Genbecev-Krtolica, O., Mavrogianis, P., Ferrer, K., Kruessel, J., et al. (2007). Decidual stromal cell response to paracrine signals from the trophoblast: amplification of immune and angiogenic modulators. Biol Reprod 76, 102–117.

17. Hiby, S.E., Walker, J.J., O’Shaughnessy K, M., Redman, C.W., Carrington, M., Trowsdale, J., and Moffett, A. (2004). Combinations of maternal KIR and fetal HLA-C genes influence the risk of preeclampsia and reproductive success. J Exp Med 200, 957–965.

18. Hunt, J.S., Petroff, M.G., McIntire, R.H., and Ober, C. (2005). HLA-G and immune tolerance in pregnancy. FASEB J 19, 681–693.

19. Irwin, J.C., and Giudice, L.C. (1998). Insulin-like growth factor binding protein-1 binds to placental cytotrophoblast alpha5beta1 integrin and inhibits cytotrophoblast invasion into decidualized endometrial stromal cultures. Growth Horm IGF Res 8, 21–31.

20. James, J.L., Stone, P.R., and Chamley, L.W. (2005). Cytotrophoblast differentiation in the first trimester of pregnancy: evidence for separate progenitors of extravillous trophoblasts and syncytiotrophoblast. Reproduction 130, 95–103.

21. Jin, S., Guerrero-Juarez, C.F., Zhang, L., Chang, I., Ramos, R., Kuan, C.H., Myung, P., Plikus, M.V., and Nie, Q. (2021). Inference and analysis of cell-cell communication using CellChat. Nat Commun 12, 1088.

22. Kamimoto, K., Stringa, B., Hoffmann, C.M., Jindal, K., Solnica-Krezel, L., and Morris, S.A. (2023). Dissecting cell identity via network inference and in silico gene perturbation. Nature 614, 742–751.

23. Kashiwagi, H., Shiraga, M., Kato, H., Kamae, T., Yamamoto, N., Tadokoro, S., Kurata, Y., Tomiyama, Y., and Kanakura, Y. (2005). Negative regulation of platelet function by a secreted cell repulsive protein, semaphorin 3A. Blood 106, 913–921.

24. Keren, L., Bosse, M., Thompson, S., Risom, T., Vijayaragavan, K., McCaffrey, E., Marquez, D., Angoshtari, R., Greenwald, N.F., Fienberg, H., et al. (2019). MIBI-TOF: A multiplexed imaging platform relates cellular phenotypes and tissue structure. Sci Adv 5, eaax5851.

25. Labarrere, C.A., DiCarlo, H.L., Bammerlin, E., Hardin, J.W., Kim, Y.M., Chaemsaithong, P., Haas, D.M., Kassab, G.S., and Romero, R. (2017). Failure of physiologic transformation of spiral arteries, endothelial and trophoblast cell activation, and acute atherosis in the basal plate of the placenta. Am J Obstet Gynecol 216, 287 e281–287 e216.

26. Laisk, T., Soares, A.L.G., Ferreira, T., Painter, J.N., Censin, J.C., Laber, S., Bacelis, J., Chen, C.Y., Lepamets, M., Lin, K., et al. (2020). The genetic architecture of sporadic and multiple consecutive miscarriage. Nat Commun 11, 5980.

27. Lee, A.C., Blencowe, H., and Lawn, J.E. (2019). Small babies, big numbers: global estimates of preterm birth. Lancet Glob Health 7, e2–e3.

28. Lee, C.Q.E., Turco, M.Y., Gardner, L., Simons, B.D., Hemberger, M., and Moffett, A. (2018). Integrin alpha2 marks a niche of trophoblast progenitor cells in first trimester human placenta. Development 145.

29. Liu, Y., Fan, X., Wang, R., Lu, X., Dang, Y.L., Wang, H., Lin, H.Y., Zhu, C., Ge, H., Cross, J.C., et al. (2018). Single-cell RNA-seq reveals the diversity of trophoblast subtypes and patterns of differentiation in the human placenta. Cell Res 28, 819–832.

30. Lokossou, A.G., Toudic, C., and Barbeau, B. (2014). Implication of human endogenous retrovirus envelope proteins in placental functions. Viruses 6, 4609–4627.

31. Marsh, B., Zhou, Y., Kapidzic, M., Fisher, S., and Blelloch, R. (2022). Regionally distinct trophoblast regulate barrier function and invasion in the human placenta. Elife 11.

32. Miranda, A.L., Kourdova, L.T., Racca, A.C., Cruz Del Puerto, M., Rojas, M.L., Marques, A.L.X., Silva, E.C.O., Fonseca, E.J.S., Gazzoni, Y., Gruppi, A., et al. (2022). Kruppel-like factor 6 participates in extravillous trophoblast cell differentiation and its expression is reduced in abnormally invasive placenta. FEBS Lett 596, 1700–1719.

33. Oliver-Williams, C., Fleming, M., Wood, A.M., and Smith, G. (2015). Previous miscarriage and the subsequent risk of preterm birth in Scotland, 1980-2008: a historical cohort study. BJOG 122, 1525-1534.

34. Omori, K., and Kotera, J. (2007). Overview of PDEs and their regulation. Circ Res 100, 309–327.

35. Ozeki, A., Oogaki, Y., Henmi, Y., Karasawa, T., Takahashi, M., Takahashi, H., Ohkuchi, A., and Shirasuna, K. (2022). Elevated S100A9 in preeclampsia induces soluble endoglin and IL-1beta secretion and hypertension via the NLRP3 inflammasome. J Hypertens 40, 84–93.

36. Pawlak, J.B., Balint, L., Lim, L., Ma, W., Davis, R.B., Benyo, Z., Soares, M.J., Oliver, G., Kahn, M.L., Jakus, Z., et al. (2019). Lymphatic mimicry in maternal endothelial cells promotes placental spiral artery remodeling. J Clin Invest 129, 4912–4921.

37. Pique-Regi, R., Romero, R., Tarca, A.L., Sendler, E.D., Xu, Y., Garcia-Flores, V., Leng, Y., Luca, F., Hassan, S.S., and Gomez-Lopez, N. (2019). Single cell transcriptional signatures of the human placenta in term and preterm parturition. Elife 8.

38. Quenby, S., Gallos, I.D., Dhillon-Smith, R.K., Podesek, M., Stephenson, M.D., Fisher, J., Brosens, J.J., Brewin, J., Ramhorst, R., Lucas, E.S., et al. (2021). Miscarriage matters: the epidemiological, physical, psychological, and economic costs of early pregnancy loss. Lancet 397, 1658–1667.

39. Rana, S., Lemoine, E., Granger, J.P., and Karumanchi, S.A. (2019). Preeclampsia: Pathophysiology, Challenges, and Perspectives. Circ Res 124, 1094–1112.

40. Red-Horse, K., Zhou, Y., Genbacev, O., Prakobphol, A., Foulk, R., McMaster, M., and Fisher, S.J. (2004). Trophoblast differentiation during embryo implantation and formation of the maternal-fetal interface. J Clin Invest 114, 744–754.

41. Rozenblatt-Rosen, O., Regev, A., Oberdoerffer, P., Nawy, T., Hupalowska, A., Rood, J.E., Ashenberg, O., Cerami, E., Coffey, R.J., Demir, E., et al. (2020). The Human Tumor Atlas Network: Charting Tumor Transitions across Space and Time at Single-Cell Resolution. Cell 181, 236–249.

42. Ryckman, C., Vandal, K., Rouleau, P., Talbot, M., and Tessier, P.A. (2003). Proinflammatory activities of S100: proteins S100A8, S100A9, and S100A8/A9 induce neutrophil chemotaxis and adhesion. J Immunol 170, 3233-3242.

43. Sato, Y., Higuchi, T., Yoshioka, S., Tatsumi, K., Fujiwara, H., and Fujii, S. (2003). Trophoblasts acquire a chemokine receptor, CCR1, as they differentiate towards invasive phenotype. Development 130, 5519-5532.

44. Schaub, M.A., Boyle, A.P., Kundaje, A., Batzoglou, S., and Snyder, M. (2012). Linking disease associations with regulatory information in the human genome. Genome Res 22, 1748–1759.

45. Schep, A.N., Wu, B., Buenrostro, J.D., and Greenleaf, W.J. (2017). chromVAR: inferring transcription-factor-associated accessibility from single-cell epigenomic data. Nat Methods 14, 975–978.

46. Schupp, J.C., Adams, T.S., Cosme, C., Jr., Raredon, M.S.B., Yuan, Y., Omote, N., Poli, S., Chioccioli, M., Rose, K.A., Manning, E.P., et al. (2021). Integrated Single-Cell Atlas of Endothelial Cells of the Human Lung. Circulation 144, 286–302.

47. Schurch, C.M., Bhate, S.S., Barlow, G.L., Phillips, D.J., Noti, L., Zlobec, I., Chu, P., Black, S., Demeter, J., McIlwain, D.R., et al. (2020). Coordinated Cellular Neighborhoods Orchestrate Antitumoral Immunity at the Colorectal Cancer Invasive Front. Cell 182, 1341–1359 e1319.

48. Setty, M., Kiseliovas, V., Levine, J., Gayoso, A., Mazutis, L., and Pe’er, D. (2019). Characterization of cell fate probabilities in single-cell data with Palantir. Nat Biotechnol 37, 451–460.

49. Shen, L., Diao, Z., Sun, H.X., Yan, G.J., Wang, Z., Li, R.T., Dai, Y., Wang, J., Li, J., Ding, H., et al. (2017). Up-regulation of CD81 inhibits cytotrophoblast invasion and mediates maternal endothelial cell dysfunction in preeclampsia. Proc Natl Acad Sci U S A 114, 1940–1945.

50. Staff, A.C., Fjeldstad, H.E., Fosheim, I.K., Moe, K., Turowski, G., Johnsen, G.M., Alnaes-Katjavivi, P., and Sugulle, M. (2022). Failure of physiological transformation and spiral artery atherosis: their roles in preeclampsia. Am J Obstet Gynecol 226, S895–S906.

51. Steinthorsdottir, V., McGinnis, R., Williams, N.O., Stefansdottir, L., Thorleifsson, G., Shooter, S., Fadista, J., Sigurdsson, J.K., Auro, K.M., Berezina, G., et al. (2020). Genetic predisposition to hypertension is associated with preeclampsia in European and Central Asian women. Nat Commun 11, 5976.

52. Stuart, T., Srivastava, A., Madad, S., Lareau, C.A., and Satija, R. (2021). Single-cell chromatin state analysis with Signac. Nat Methods 18, 1333–1341.

53. Sudlow, C., Gallacher, J., Allen, N., Beral, V., Burton, P., Danesh, J., Downey, P., Elliott, P., Green, J., Landray, M., et al. (2015). UK biobank: an open access resource for identifying the causes of a wide range of complex diseases of middle and old age. PLoS Med 12, e1001779.

54. Suryawanshi, H., Morozov, P., Straus, A., Sahasrabudhe, N., Max, K.E.A., Garzia, A., Kustagi, M., Tuschl, T., and Williams, Z. (2018). A single-cell survey of the human first-trimester placenta and decidua. Sci Adv 4, eaau4788.

55. Swingle, H.M., Colaizy, T.T., Zimmerman, M.B., and Morriss, F.H., Jr. (2009). Abortion and the risk of subsequent preterm birth: a systematic review with meta-analyses. J Reprod Med 54, 95–108.

56. Tsang, J.C.H., Vong, J.S.L., Ji, L., Poon, L.C.Y., Jiang, P., Lui, K.O., Ni, Y.B., To, K.F., Cheng, Y.K.Y., Chiu, R.W.K., et al. (2017). Integrative single-cell and cell-free plasma RNA transcriptomics elucidates placental cellular dynamics. Proc Natl Acad Sci U S A 114, E7786–E7795.

57. Varberg, K.M., Iqbal, K., Muto, M., Simon, M.E., Scott, R.L., Kozai, K., Choudhury, R.H., Aplin, J.D., Biswell, R., Gibson, M., et al. (2021). ASCL2 reciprocally controls key trophoblast lineage decisions during hemochorial placenta development. Proc Natl Acad Sci U S A 118.

58. Vento-Tormo, R., Efremova, M., Botting, R.A., Turco, M.Y., Vento-Tormo, M., Meyer, K.B., Park, J.E., Stephenson, E., Polanski, K., Goncalves, A., et al. (2018). Single-cell reconstruction of the early maternal-fetal interface in humans. Nature 563, 347–353.

59. Vinketova, K., Mourdjeva, M., and Oreshkova, T. (2016). Human Decidual Stromal Cells as a Component of the Implantation Niche and a Modulator of Maternal Immunity. J Pregnancy 2016, 8689436.

60. Weeraratna, A.T., Jiang, Y., Hostetter, G., Rosenblatt, K., Duray, P., Bittner, M., and Trent, J.M. (2002). Wnt5a signaling directly affects cell motility and invasion of metastatic melanoma. Cancer Cell 1, 279–288.

61. Yu, F., Cato, L.D., Weng, C., Liggett, L.A., Jeon, S., Xu, K., Chiang, C.W.K., Wiemels, J.L., Weissman, J.S., de Smith, A.J., et al. (2022). Variant to function mapping at single-cell resolution through network propagation. bioRxiv.

62. Zhang, G., Feenstra, B., Bacelis, J., Liu, X., Muglia, L.M., Juodakis, J., Miller, D.E., Litterman, N., Jiang, P.P., Russell, L., et al. (2017). Genetic Associations with Gestational Duration and Spontaneous Preterm Birth. N Engl J Med 377, 1156–1167.

63. Zhang, J., Dong, H., Wang, B., Zhu, S., and Croy, B.A. (2008). Dynamic changes occur in patterns of endometrial EFNB2/EPHB4 expression during the period of spiral arterial modification in mice. Biol Reprod 79, 450–458.

64. Zhou, Y., Fisher, S.J., Janatpour, M., Genbacev, O., Dejana, E., Wheelock, M., and Damsky, C.H. (1997). Human cytotrophoblasts adopt a vascular phenotype as they differentiate. A strategy for successful endovascular invasion? J Clin Invest 99, 2139–2151.

65. Zhou, Y., McMaster, M., Woo, K., Janatpour, M., Perry, J., Karpanen, T., Alitalo, K., Damsky, C., and Fisher, S.J. (2002). Vascular endothelial growth factor ligands and receptors that regulate human cytotrophoblast survival are dysregulated in severe preeclampsia and hemolysis, elevated liver enzymes, and low platelets syndrome. Am J Pathol 160, 1405–1423.

