## Supplementary material for "A Multiomics, Spatiotemporal, and Single Cell Atlas for Mapping Cell-Type-Specific Dysregulation at the Maternal-Fetal Interface": Supp Files.pdf

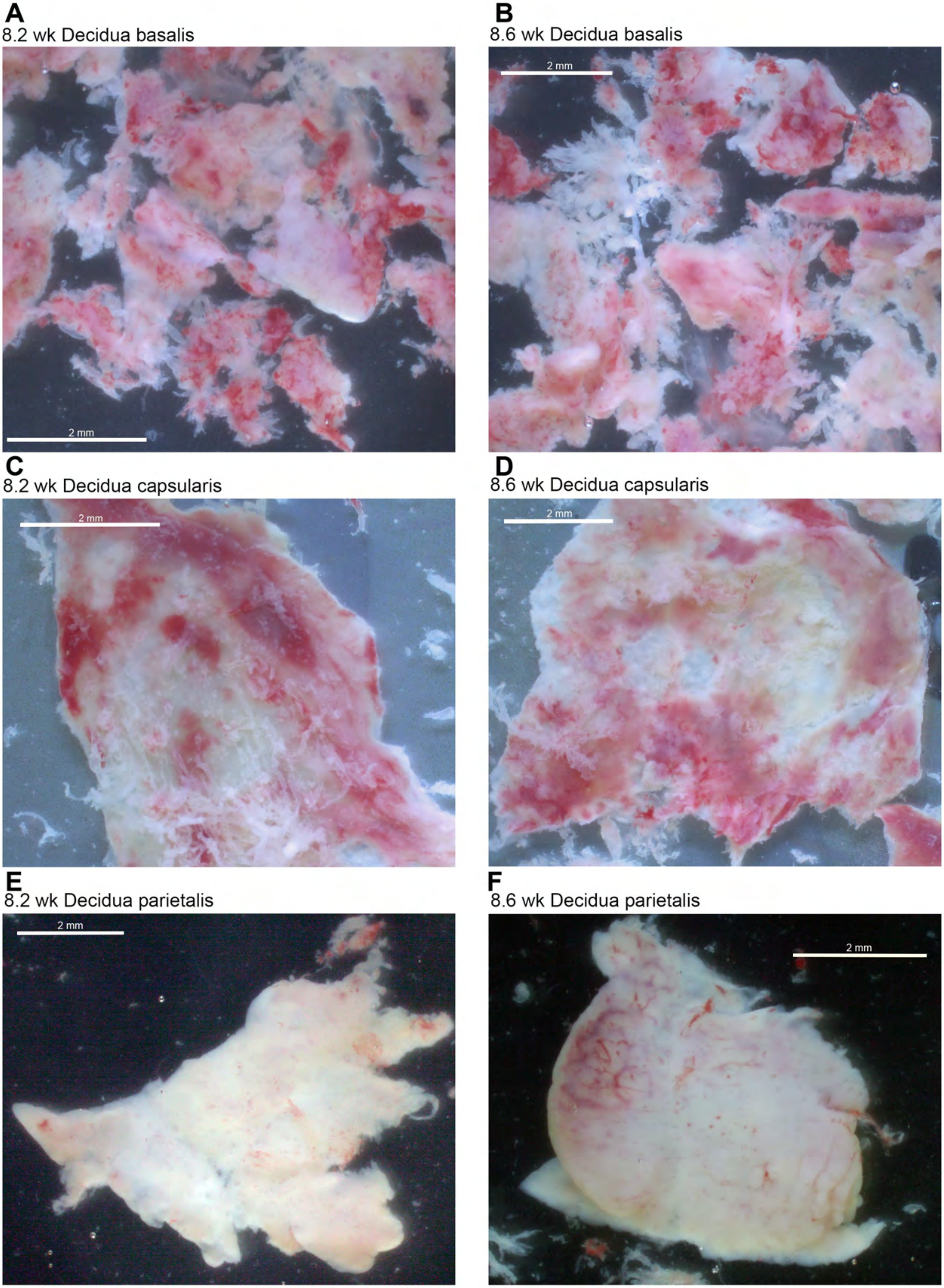
A
8.2 wk Decidua basalis
2 mm
B
8.6 wk Decidua basalis
2 mm
C
8.2 wk Decidua capsularis
2 mm
D
8.6 wk Decidua capsularis
2 mm
E
8.2 wk Decidua parietalis
2 mm
F
8.6 wk Decidua parietalis
2 mm

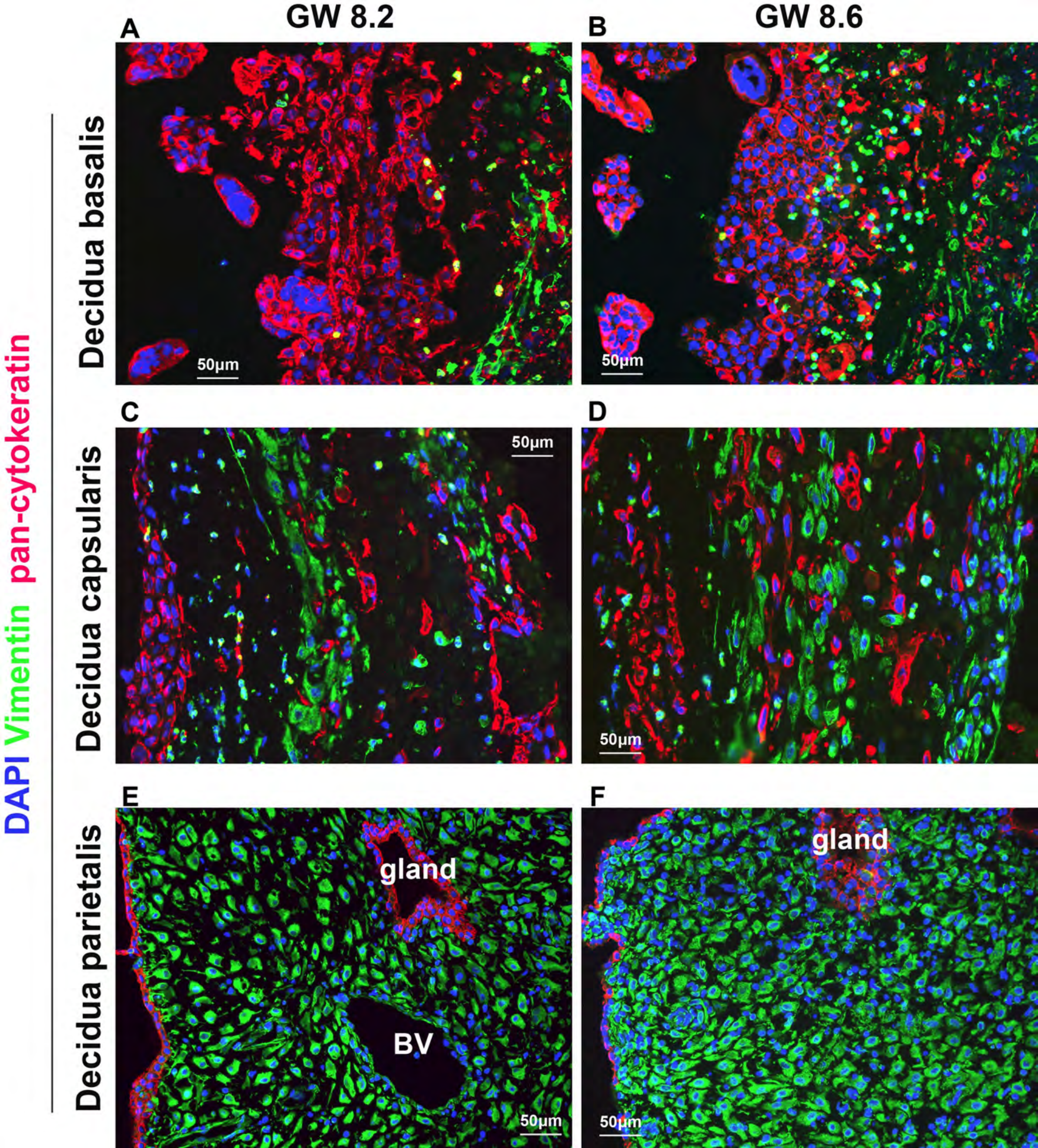
GW 8.2
GW 8.6
A
B
C
D
E
F
Decidua basalis
Decidua capsularis
Decidua parietalis
DAPI Vimentin pan-cytokeratin
50μm
gland
BV

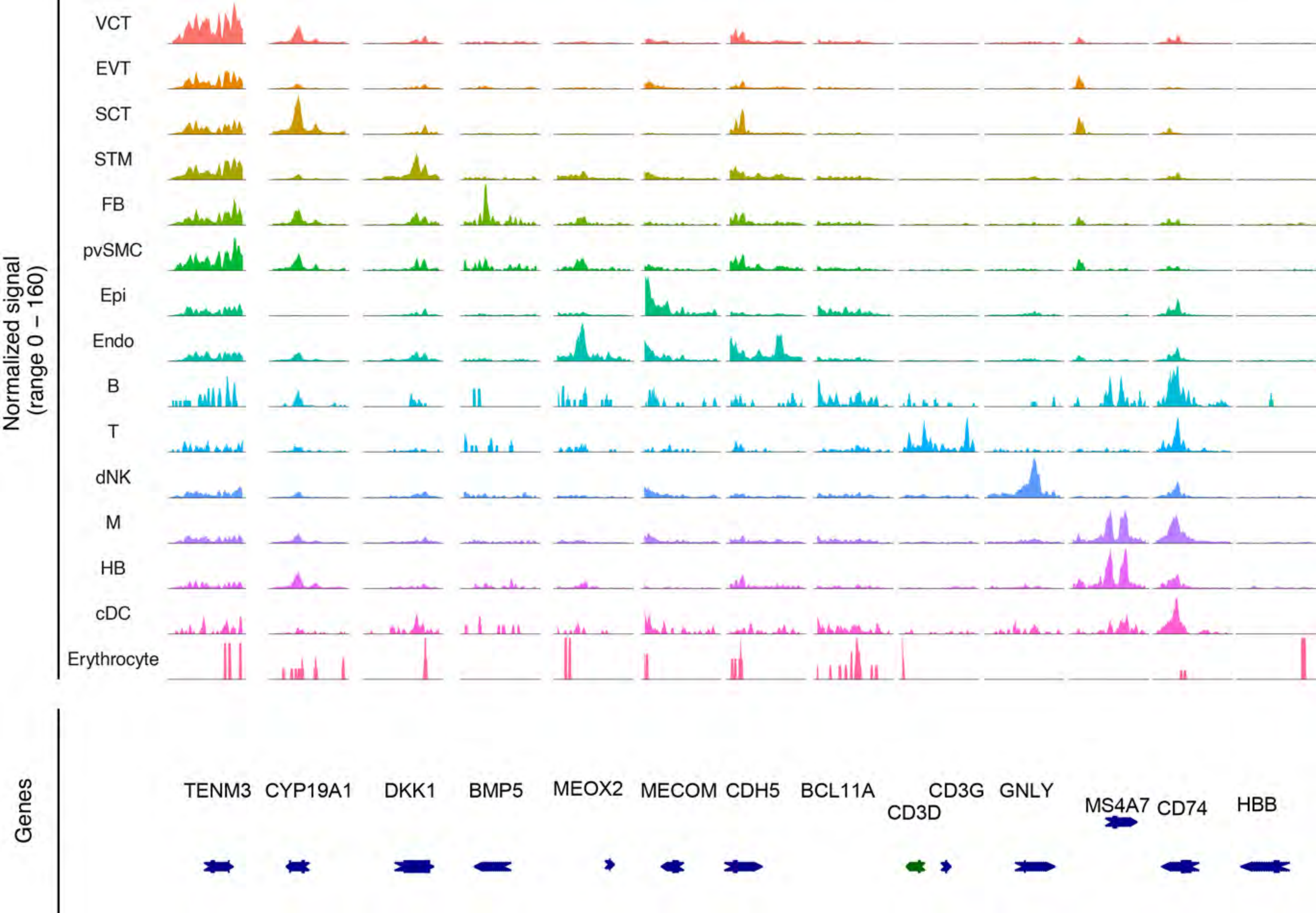

Normalized signal
(range 0 – 160)
VCT
EVT
SCT
STM
FB
pvSMC
Epi
Endo
B
T
dNK
M
HB
cDC
Erythrocyte
Genes
TENM3
CYP19A1
DKK1
BMP5
MEOX2
MECOM
CDH5
BCL11A
CD3D
CD3G
GNLY
MS4A7
CD74
HBB

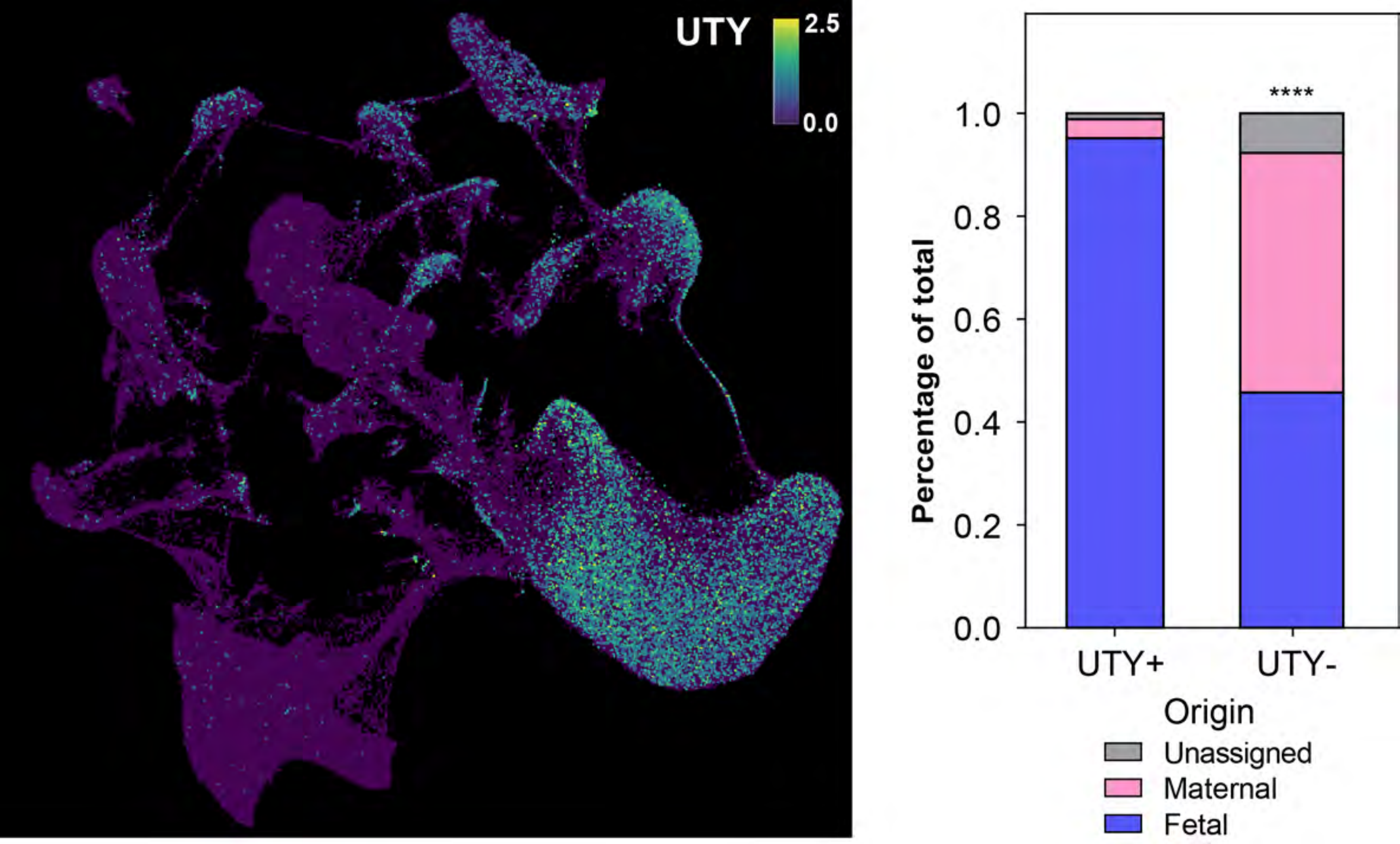
UTY
2.5
0.0
****
1.0
0.8
0.6
0.4
0.2
0.0
Percentage of total
UTY+
UTY-
Origin
Unassigned
Maternal
Fetal

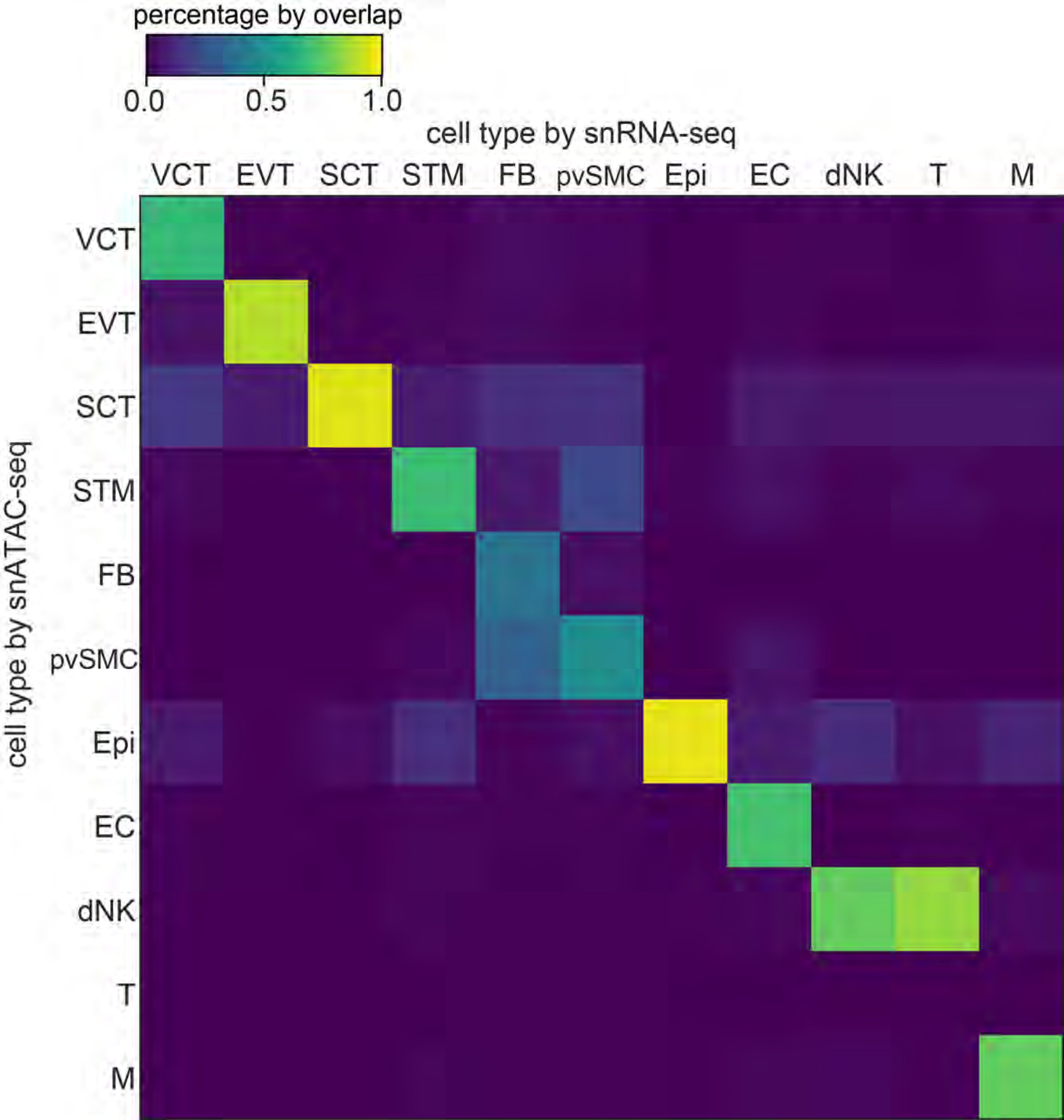

percentage by overlap
0.0
0.5
1.0
cell type by snRNA-seq
VCT
EVT
SCT
STM
FB
pvSMC
Epi
EC
dNK
T
M
cell type by snATAC-seq
VCT
EVT
SCT
STM
FB
pvSMC
Epi
EC
dNK
T
M

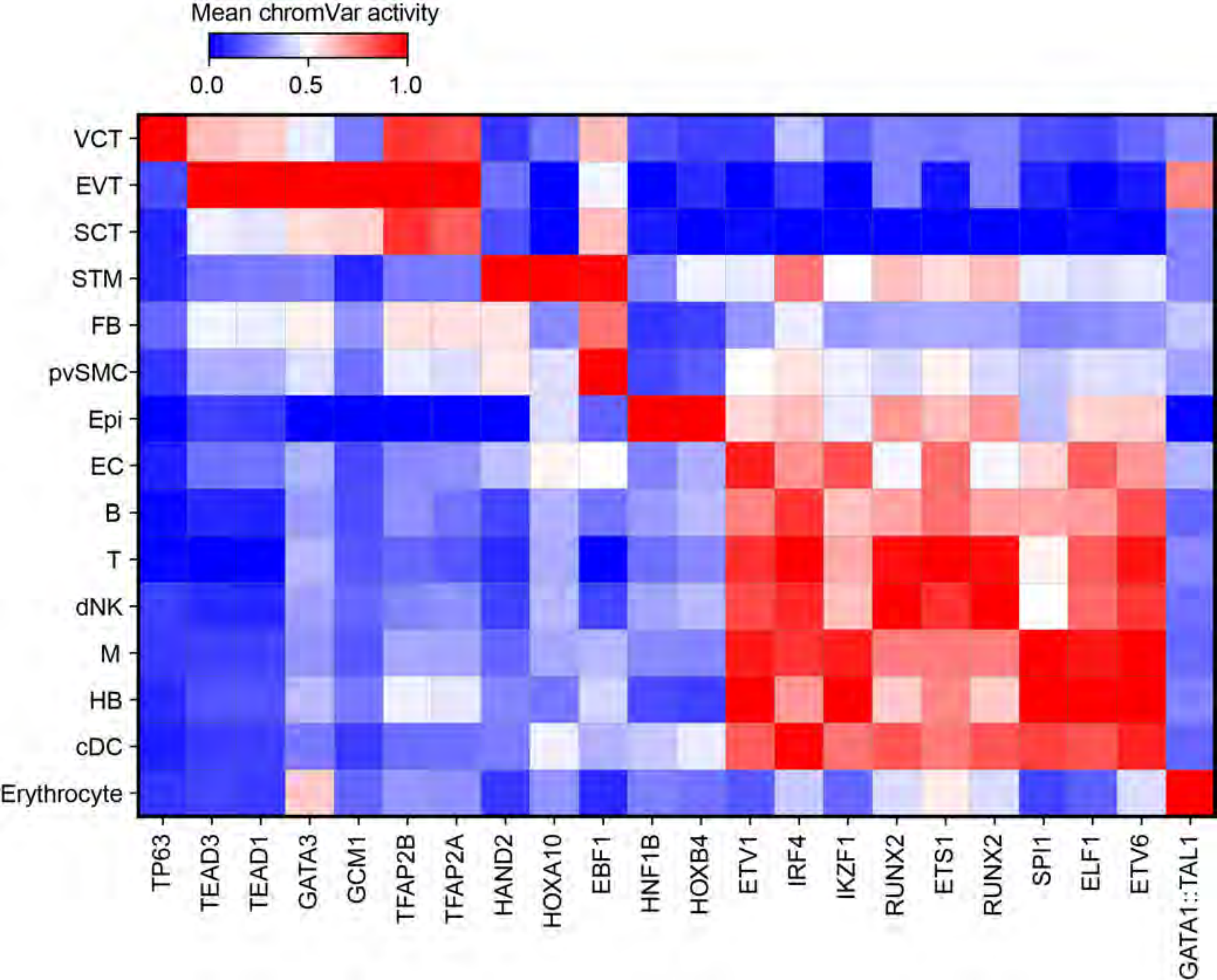
Mean chromVar activity
0.0
0.5
1.0
VCT
EVT
SCT
STM
FB
pvSMC
Epi
EC
B
T
dNK
M
HB
cDC
Erythrocyte
TP63
TEAD3
TEAD1
GATA3
GCM1
TFAP2B
TFAP2A
HAND2
HOXA10
EBF1
HNF1B
HOXB4
ETV1
IRF4
IKZF1
RUNX2
ETS1
RUNX2
SPI1
ELF1
ETV6
GATA1::TAL1

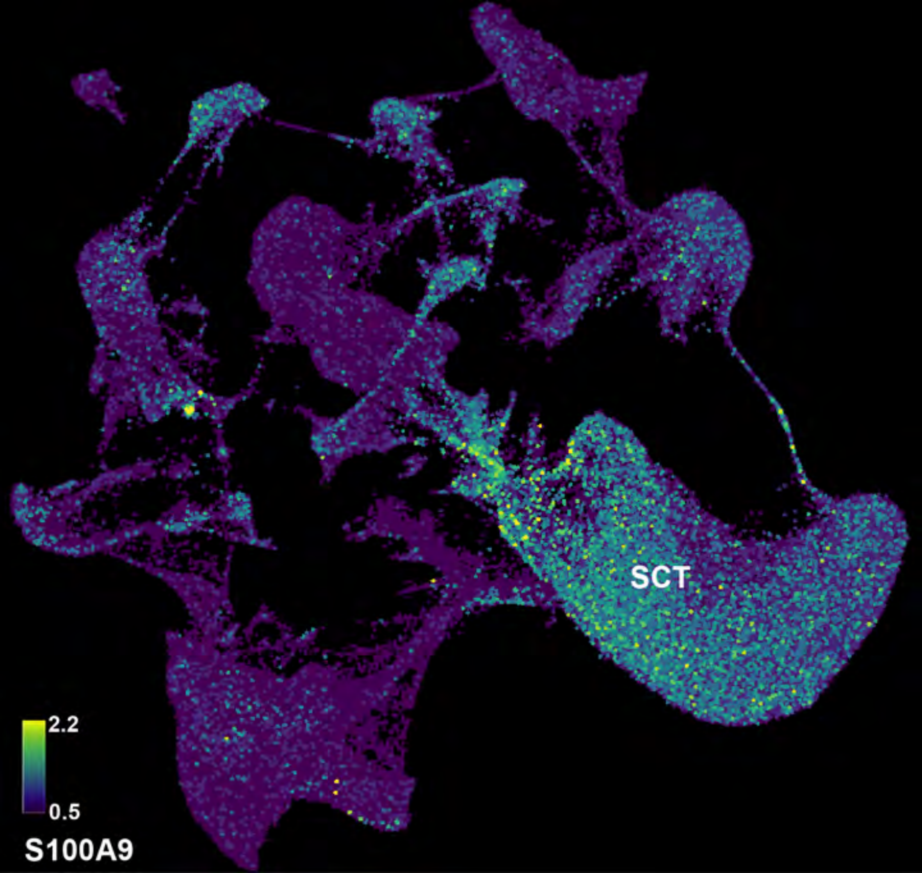
SCT
2.2
0.5
S100A9

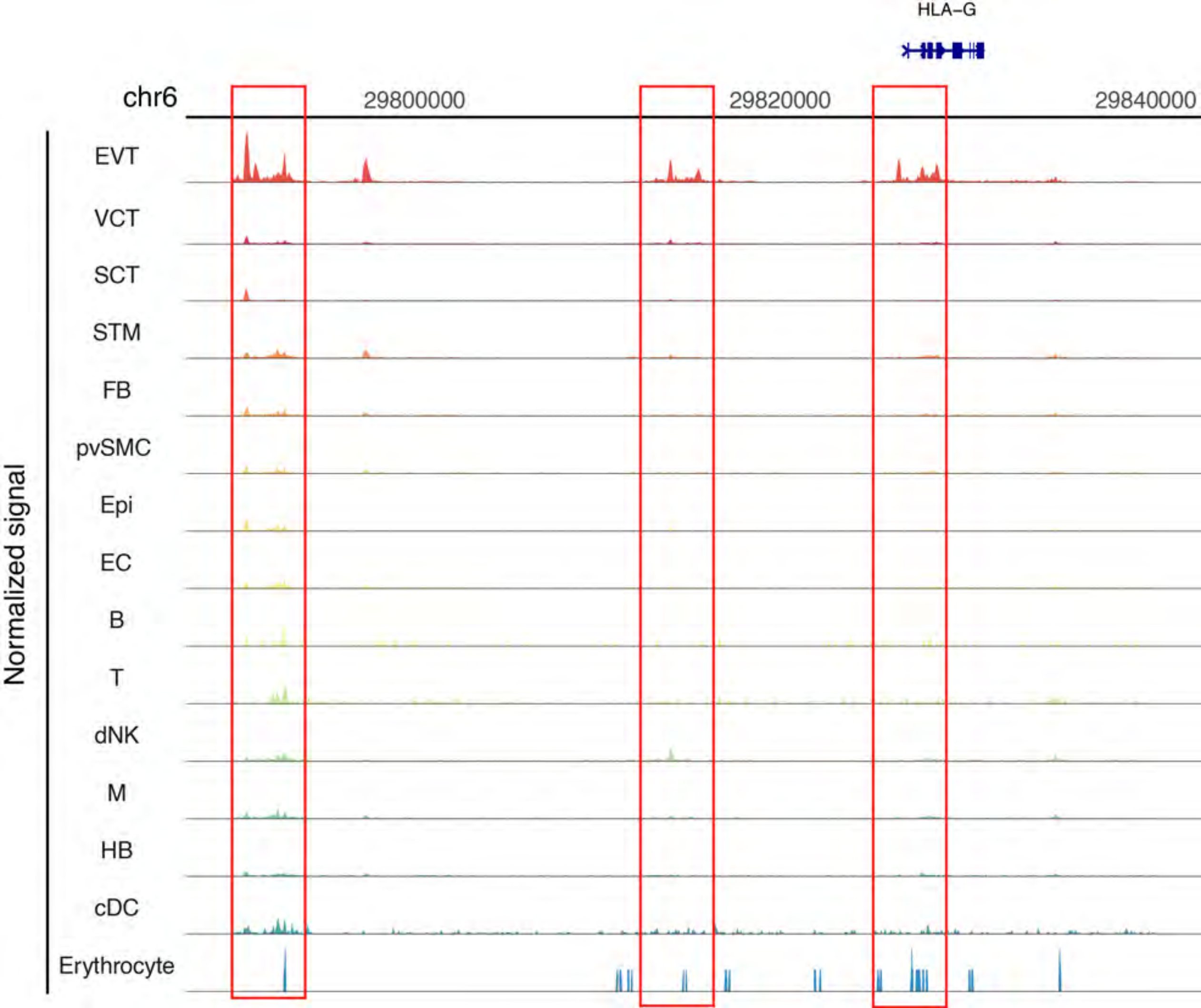
HLA−G
chr6
29800000
29820000
29840000
Normalized signal
EVT
VCT
SCT
STM
FB
pvSMC
Epi
EC
B
T
dNK
M
HB
cDC
Erythrocyte

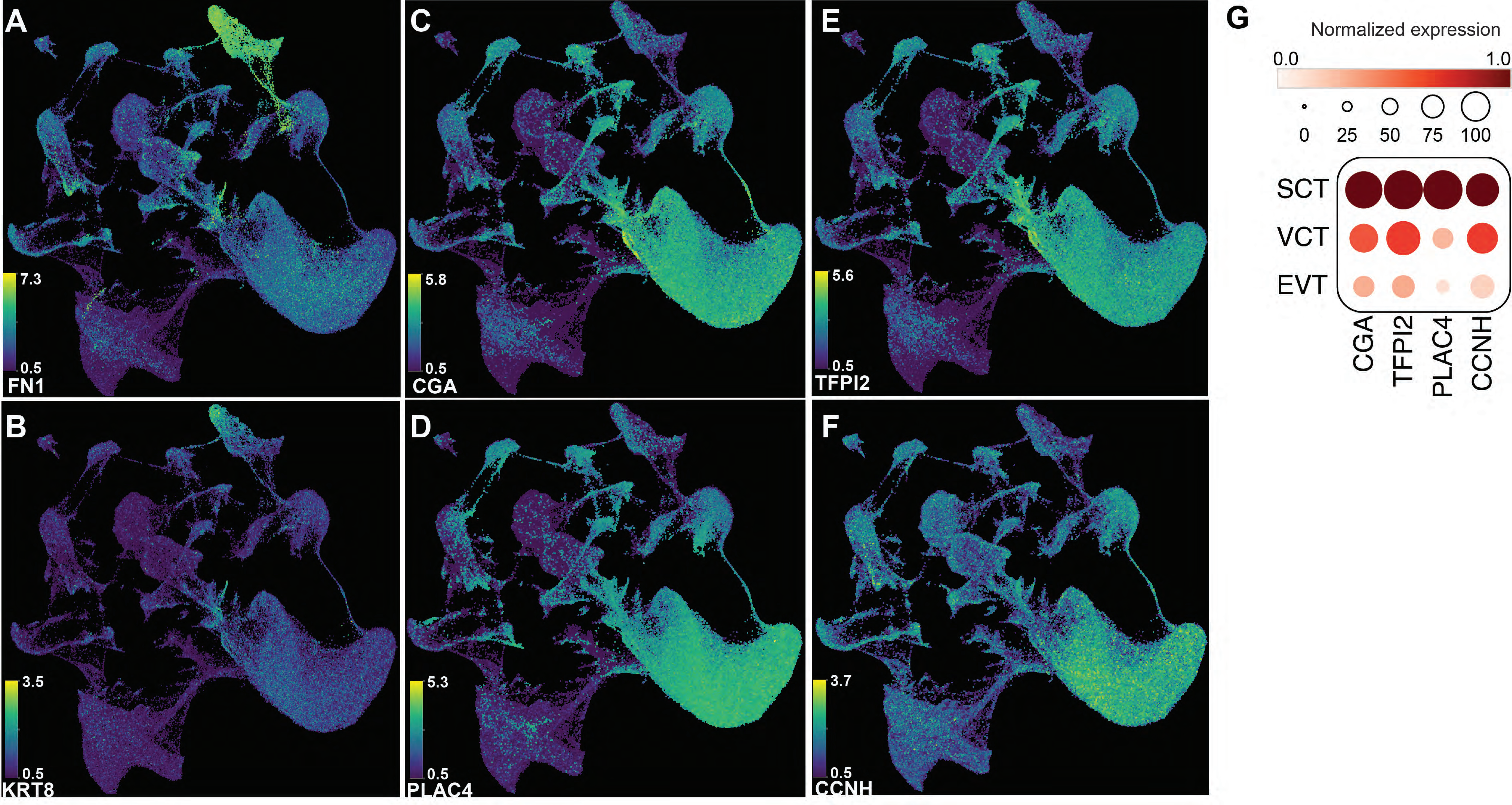

A
7.3
0.5
FN1
B
3.5
0.5
KRT8
C
5.8
0.5
CGA
D
5.3
0.5
PLAC4
E
5.6
0.5
TFPI2
F
3.7
0.5
CCNH
G
Normalized expression
0.0
1.0
0
25
50
75
100
SCT
VCT
EVT
CGA
TFPI2
PLAC4
CCNH

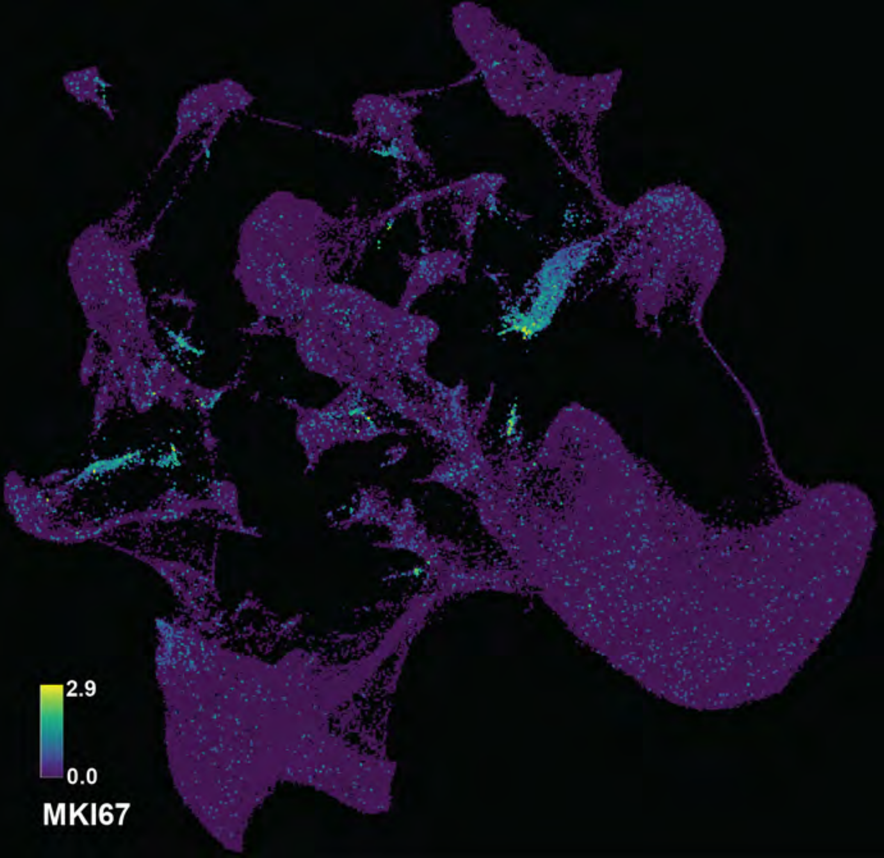
2.9
0.0
MKI67

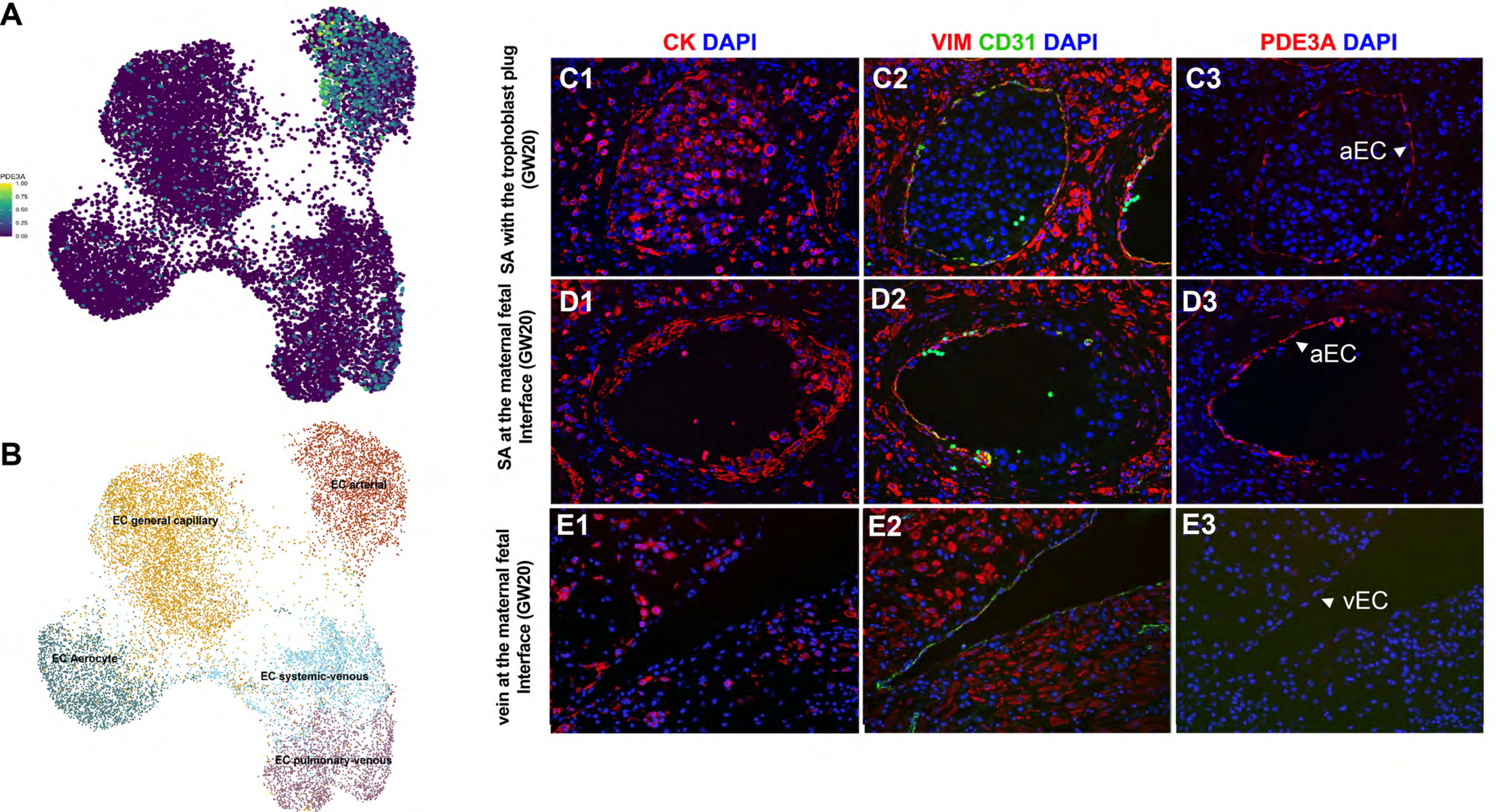
A
PDE3A
1.00
0.75
0.50
0.25
0.00
B
EC arterial
EC general capillary
EC Aerocyte
EC systemic-venous
EC pulmonary-venous
CK DAPI
VIM CD31 DAPI
PDE3A DAPI
SA with the trophoblast plug (GW20)
SA at the maternal fetal Interface (GW20)
vein at the maternal fetal Interface (GW20)
C1
C2
C3
aEC
D1
D2
D3
aEC
E1
E2
E3
vEC

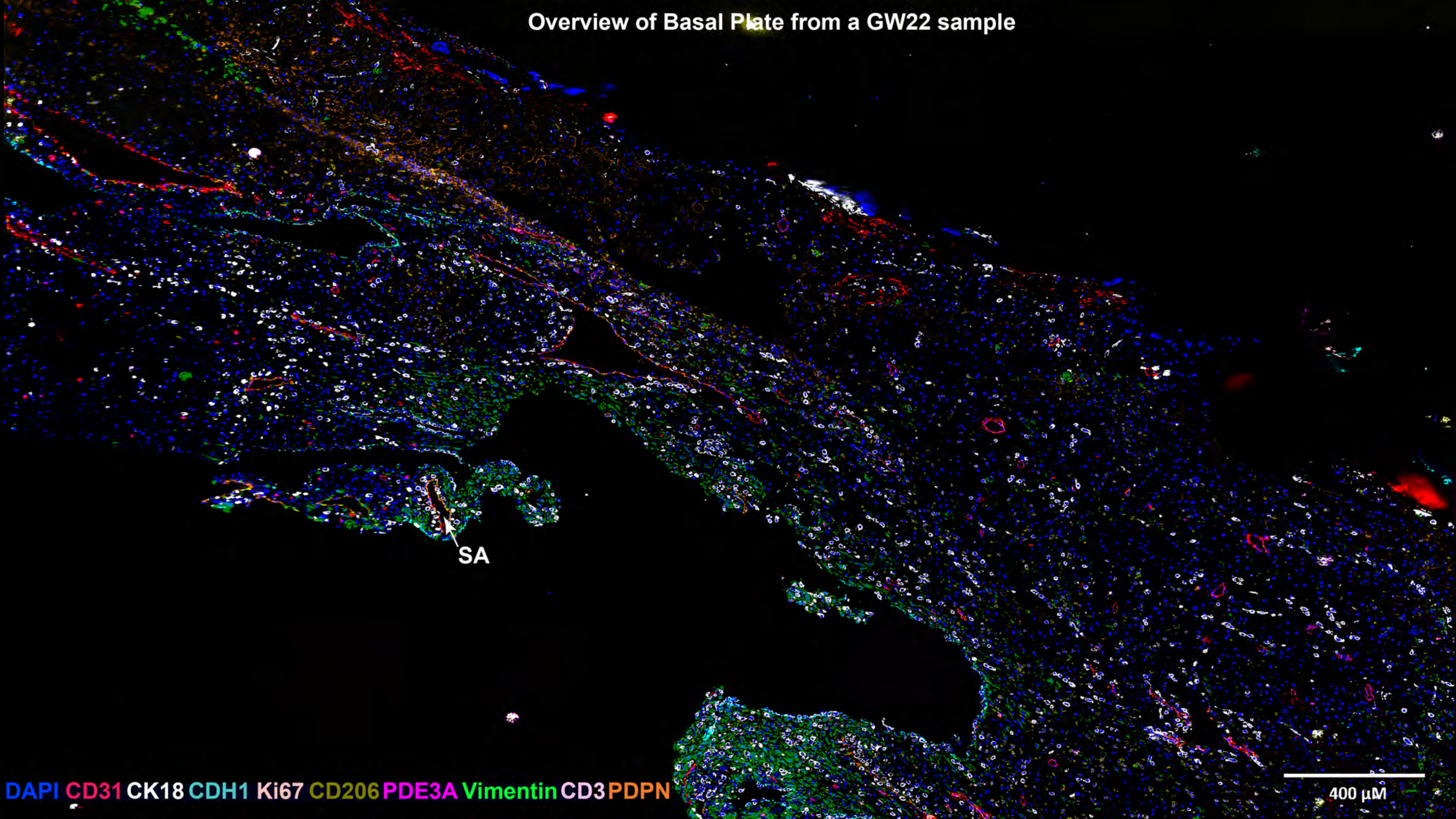
Overview of Basal Plate from a GW22 sample
SA
DAPI CD31 CK18 CDH1 Ki67 CD206 PDE3A Vimentin CD3 PDPN
400 µM

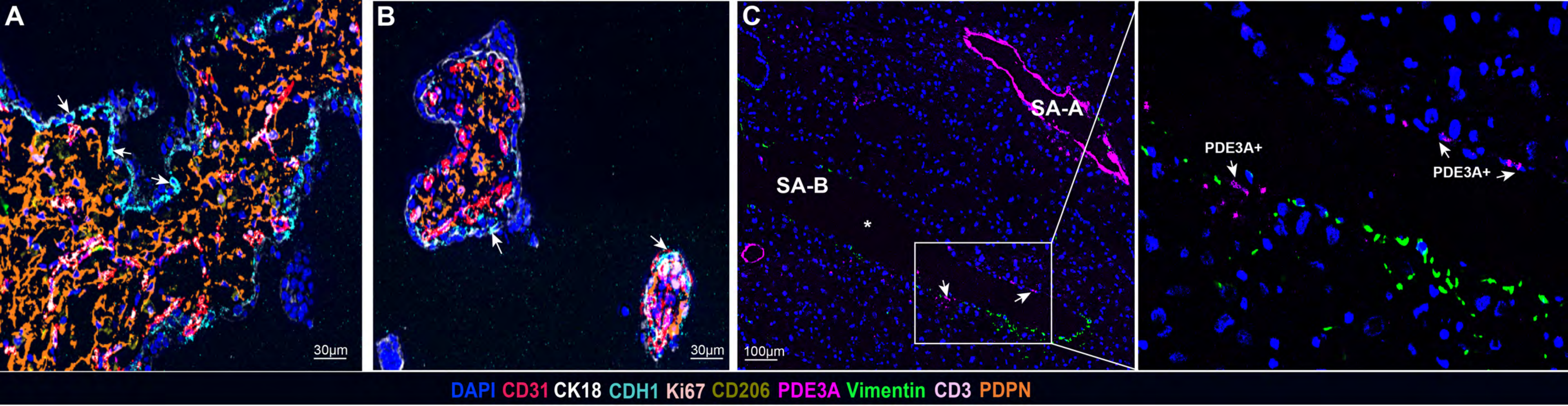
A
B
C
30µm
30µm
100µm
SA-A
SA-B
*
PDE3A+
PDE3A+
DAPI CD31 CK18 CDH1 Ki67 CD206 PDE3A Vimentin CD3 PDPN

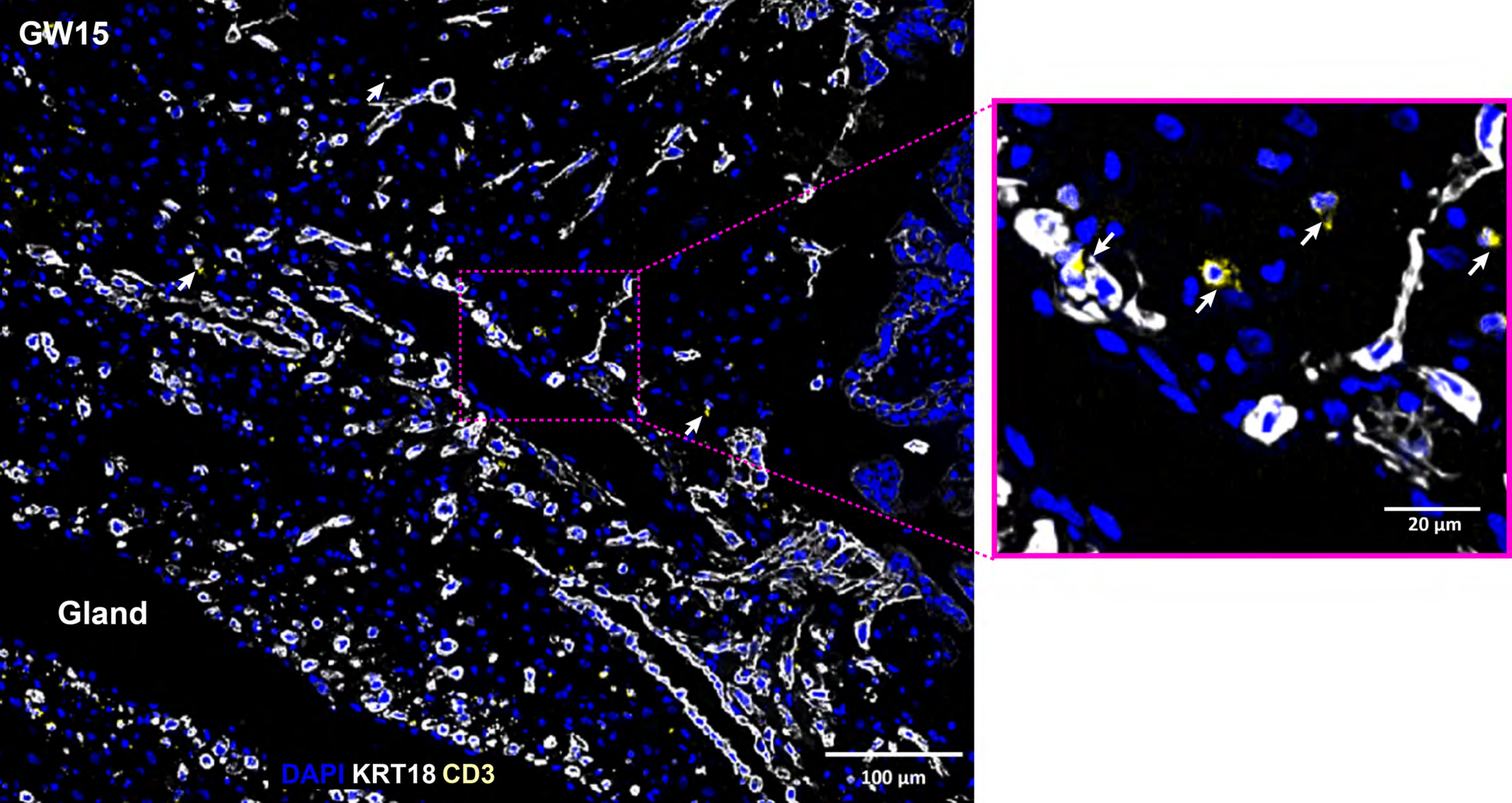

GW15
Gland
DAPI KRT18 CD3
100 µm
20 µm

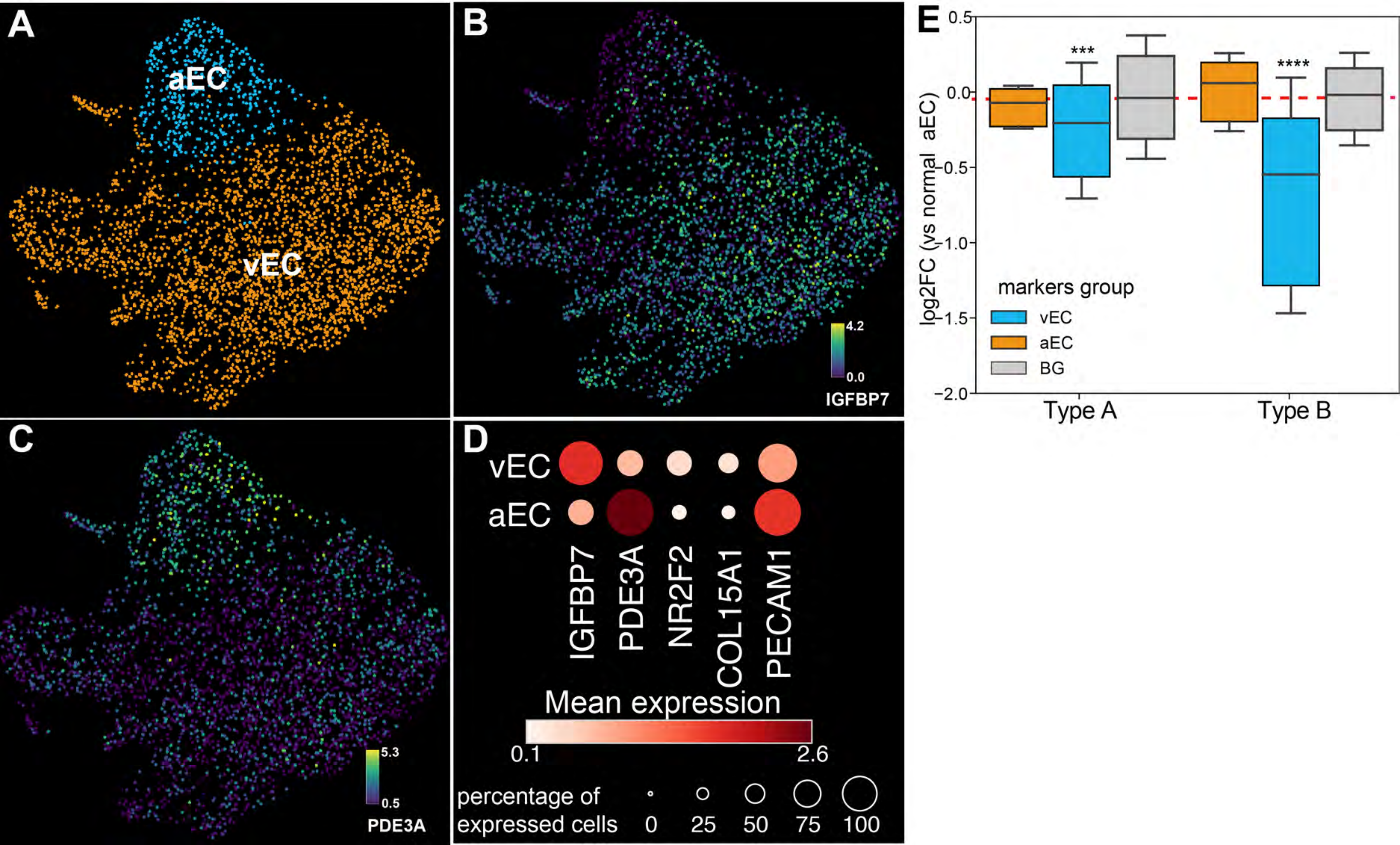
A
aEC
vEC
B
4.2
0.0
IGFBP7
C
5.3
0.5
PDE3A
D
vEC
aEC
IGFBP7
PDE3A
NR2F2
COL15A1
PECAM1
Mean expression
0.1
2.6
percentage of expressed cells
0
25
50
75
100
E
0.5
0.0
−0.5
−1.0
−1.5
−2.0
log2FC (vs normal aEC)
***
****
markers group
vEC
aEC
BG
Type A
Type B

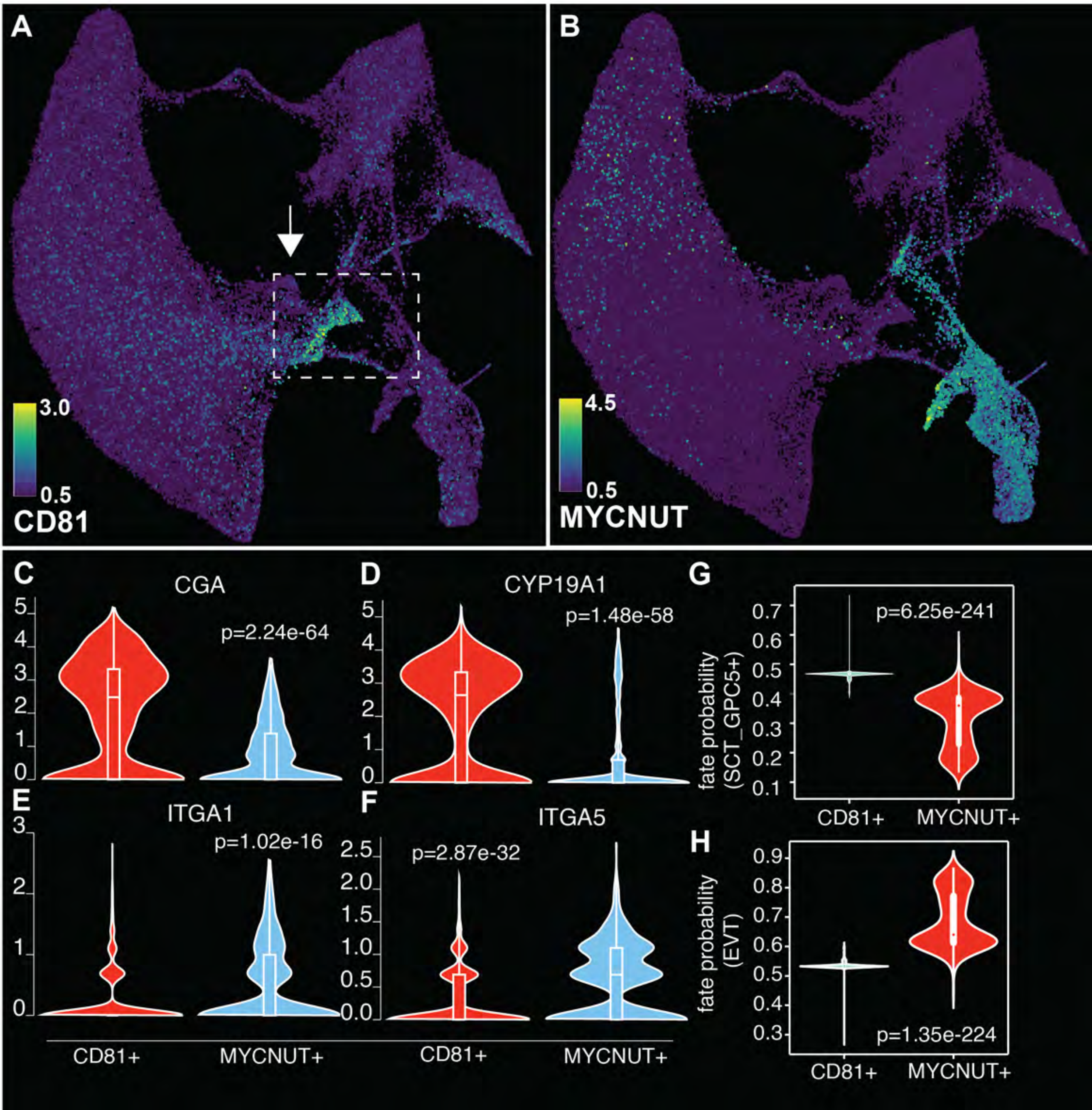
A
B
3.0
0.5
CD81
4.5
0.5
MYCNUT
C
CGA
p=2.24e-64
D
CYP19A1
p=1.48e-58
G
fate probability (SCT_GPC5+)
p=6.25e-241
E
ITGA1
p=1.02e-16
F
ITGA5
p=2.87e-32
H
fate probability (EVT)
p=1.35e-224
CD81+
MYCNUT+

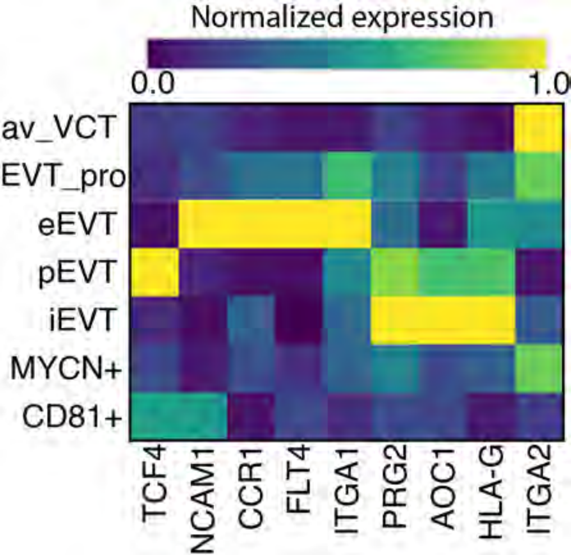
Normalized expression
0.0
1.0
av_VCT
EVT_pro
eEVT
pEVT
iEVT
MYCN+
CD81+
TCF4
NCAM1
CCR1
FLT4
ITGA1
PRG2
AOC1
HLA-G
ITGA2

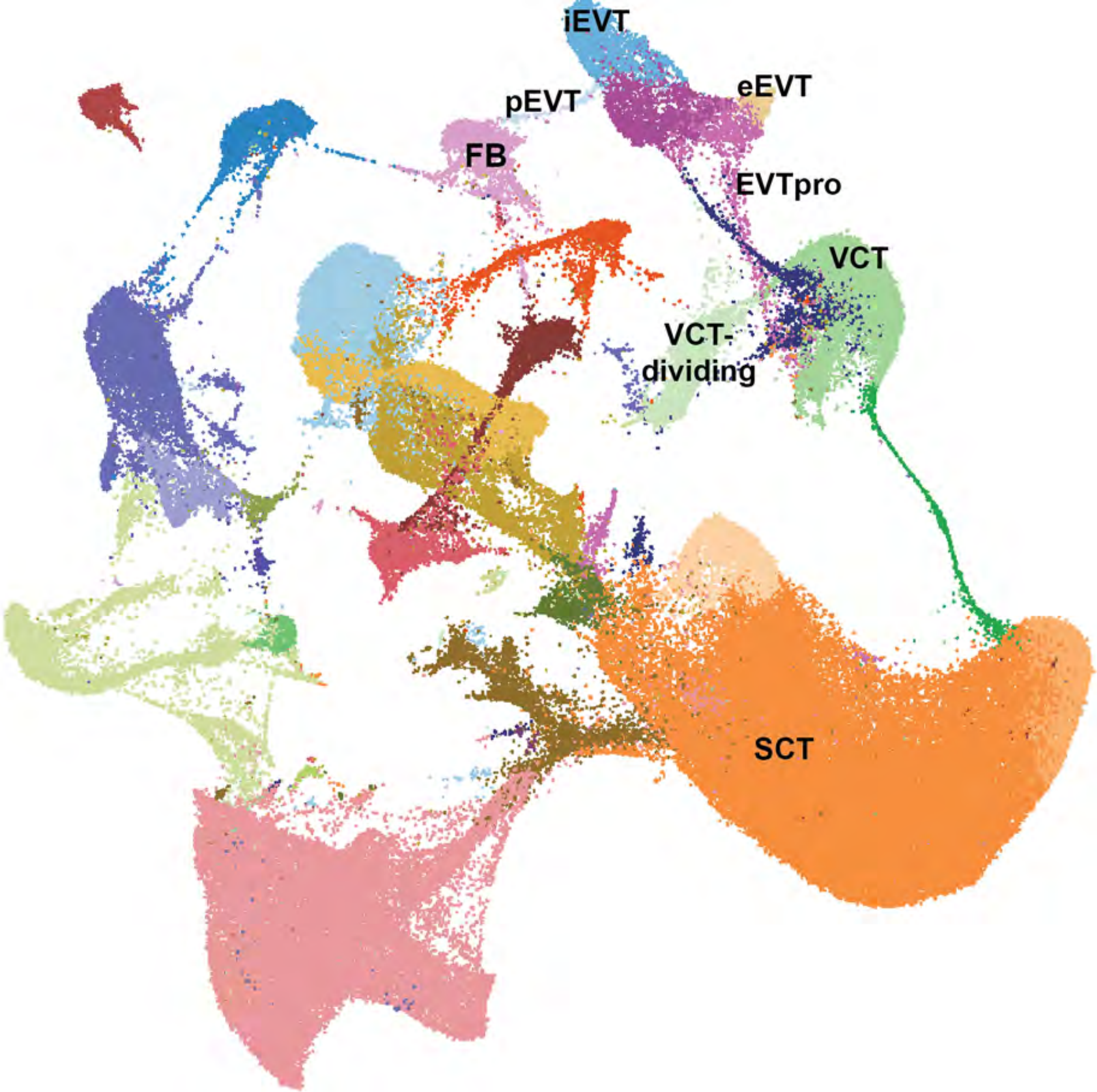
iEVT
eEVT
pEVT
FB
EVTpro
VCT
VCT-
dividing
SCT

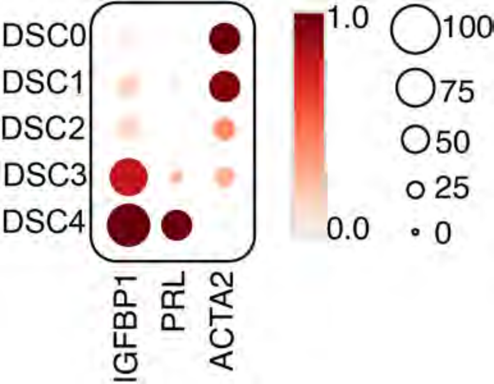
DSC0
DSC1
DSC2
DSC3
DSC4
IGFBP1
PRL
ACTA2
1.0
0.0
100
75
50
25
0

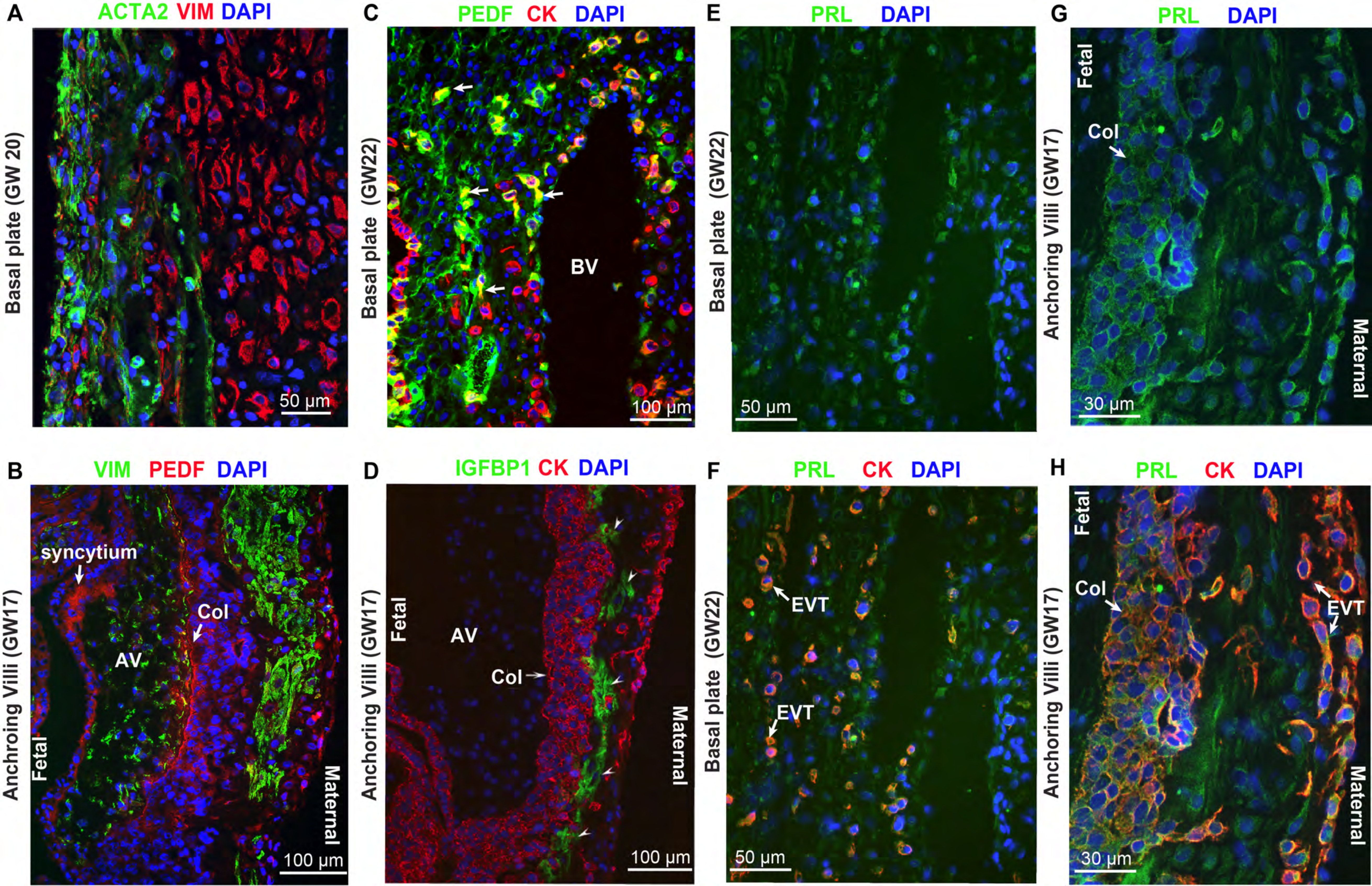

A
ACTA2 VIM DAPI
Basal plate (GW 20)
50 µm
B
VIM PEDF DAPI
Anchroing Villi (GW17)
syncytium
Col
AV
Fetal
Maternal
100 µm
C
PEDF CK DAPI
Basal plate (GW22)
BV
100 µm
D
IGFBP1 CK DAPI
Anchoring Villi (GW17)
Fetal
AV
Col
Maternal
100 µm
E
PRL DAPI
Basal plate (GW22)
50 µm
F
PRL CK DAPI
Basal plate (GW22)
EVT
EVT
50 µm
G
PRL DAPI
Anchoring Villi (GW17)
Fetal
Col
Maternal
30 µm
H
PRL CK DAPI
Anchroing Villi (GW17)
Fetal
Col
EVT
Maternal
30 µm

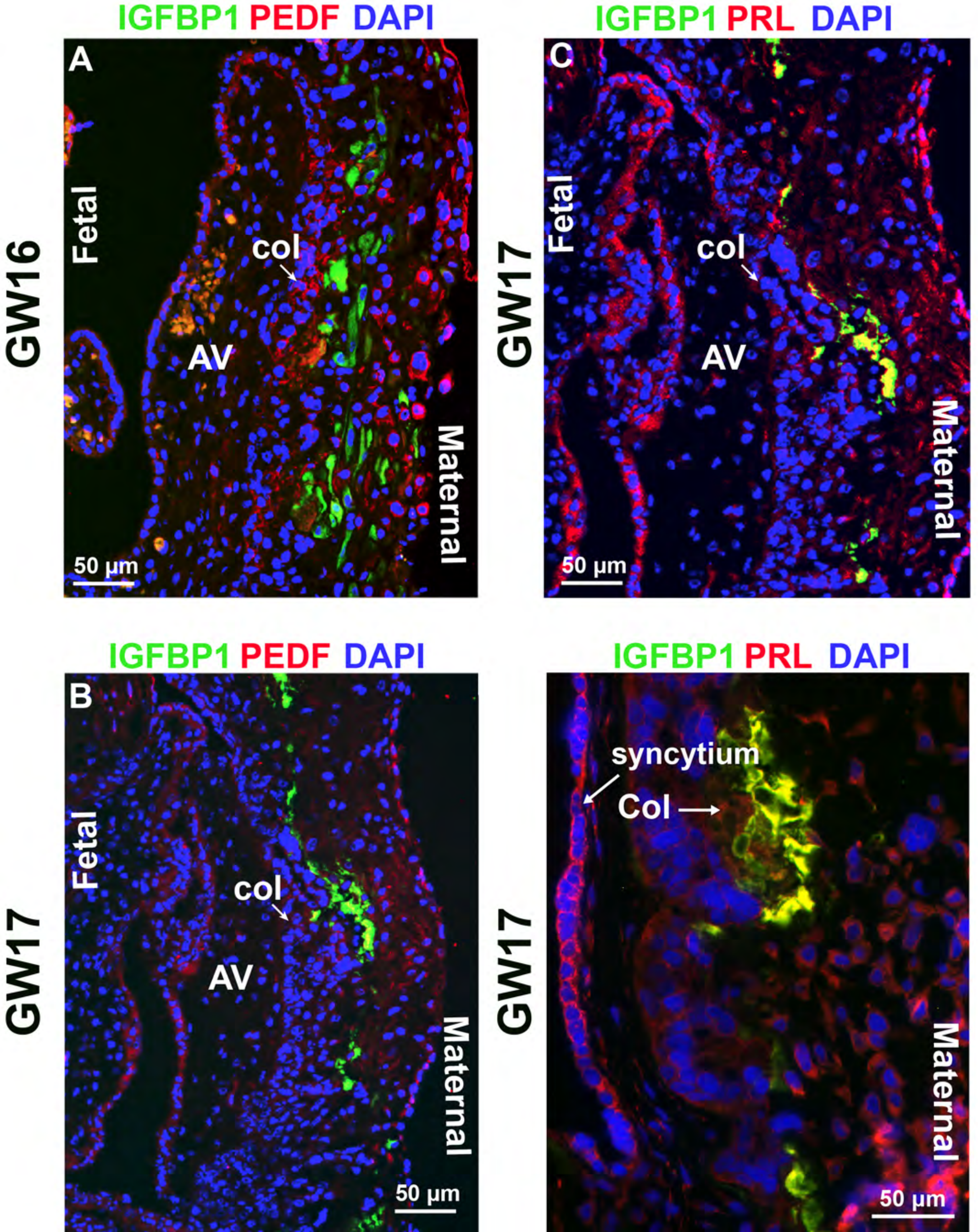
IGFBP1 PEDF DAPI
A
GW16
Fetal
col
AV
Maternal
50 µm
IGFBP1 PRL DAPI
C
GW17
Fetal
col
AV
Maternal
50 µm
IGFBP1 PEDF DAPI
B
GW17
Fetal
col
AV
Maternal
50 µm
IGFBP1 PRL DAPI
GW17
syncytium
Col
Maternal
50 µm

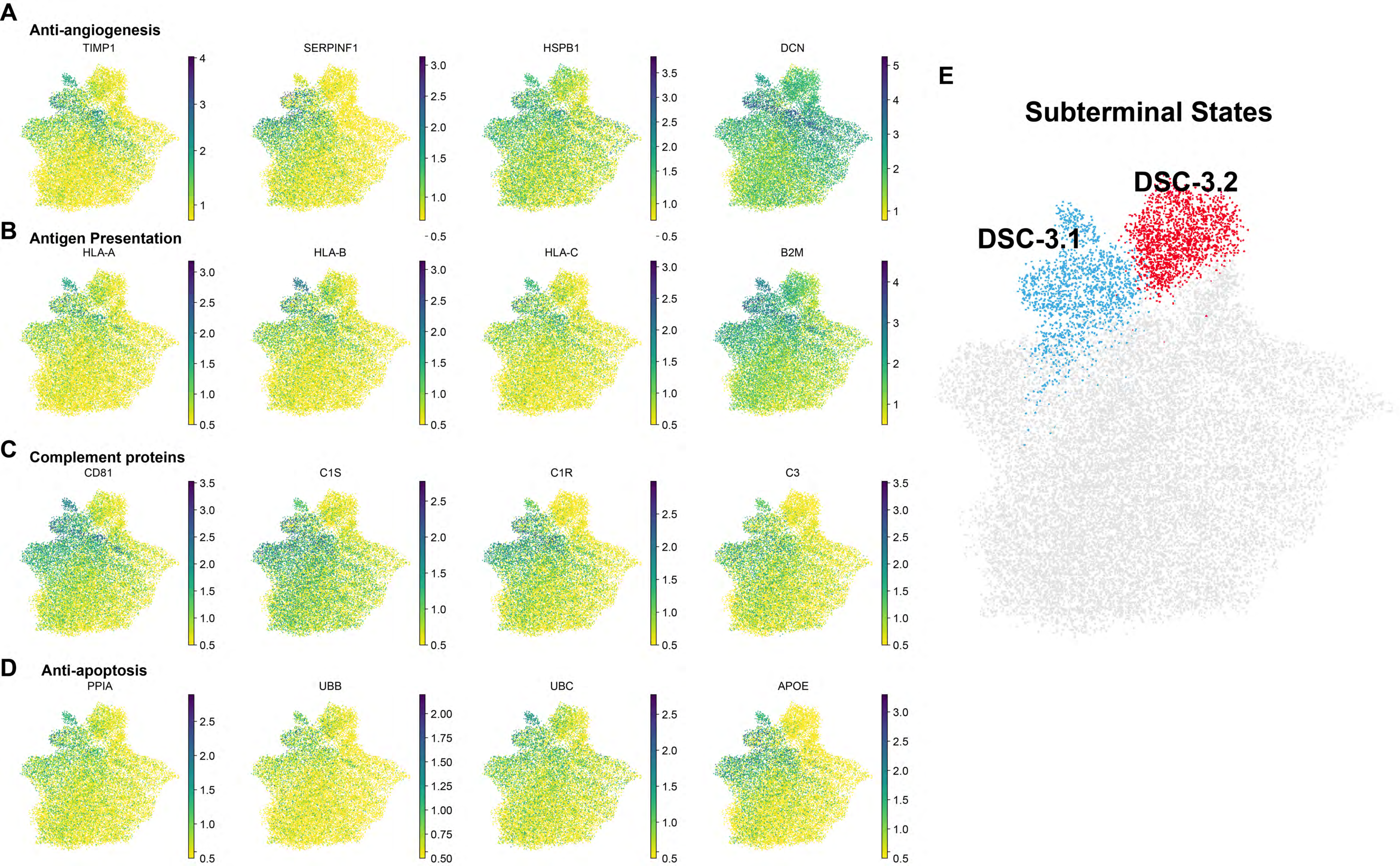
A
Anti-angiogenesis
TIMP1
SERPINF1
HSPB1
DCN
B
Antigen Presentation
HLA-A
HLA-B
HLA-C
B2M
C
Complement proteins
CD81
C1S
C1R
C3
D
Anti-apoptosis
PPIA
UBB
UBC
APOE
E
Subterminal States
DSC-3.1
DSC-3.2

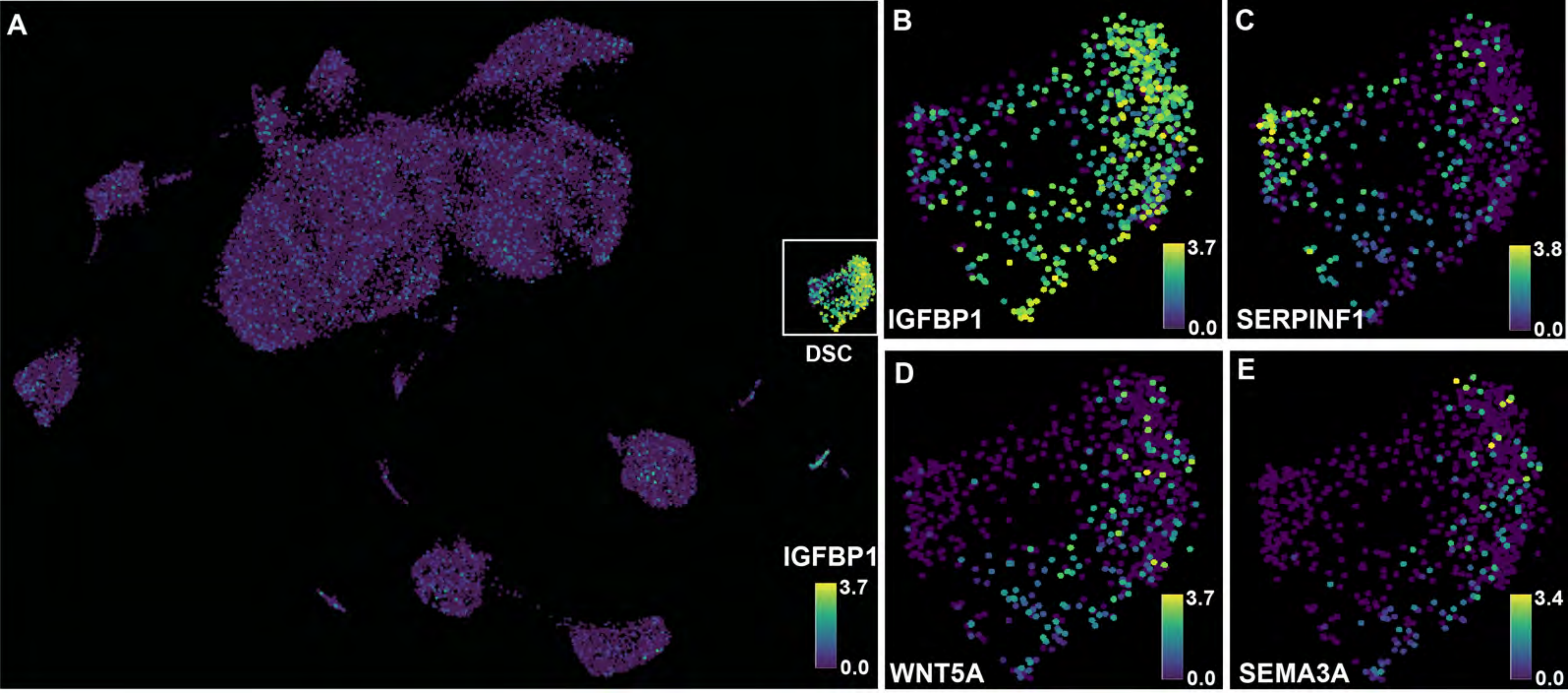
A
DSC
IGFBP1
3.7
0.0
B
IGFBP1
3.7
0.0
C
SERPINF1
3.8
0.0
D
WNT5A
3.7
0.0
E
SEMA3A
3.4
0.0

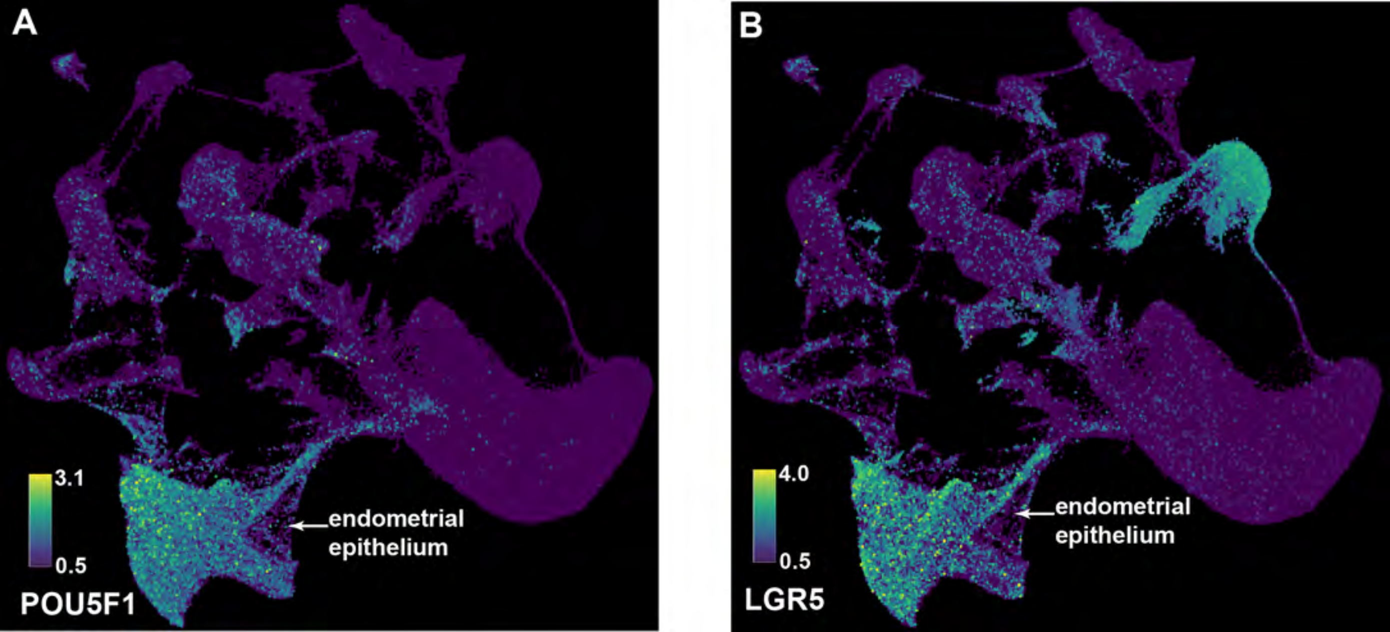
A
3.1
0.5
POU5F1
endometrial
epithelium
B
4.0
0.5
LGR5
endometrial
epithelium
