## Supplementary material for "A Multiomics, Spatiotemporal, and Single Cell Atlas for Mapping Cell-Type-Specific Dysregulation at the Maternal-Fetal Interface": Supplementary_Figure_Table_Legends.docx

**Legends for Supplementary Figures**

**Supplementary Figure 1**. The representative brightfield images showing the histological characteristics of the micro-dissected decidua basalis (**A**: GW 8.2, **B**: GW 8.6), the decidua capsularis (**C**:GW 8.2, **D**: GW 8.6), and the decidua parietalis (**E**: GW 8.2, **F**: GW 8.6). For early gestation samples, we only collected the decidua basalis samples for single cell analysis in this study.

**Supplementary Figure 2.** Immunochemical experiments using antibodies against Vimentin (green) and pan-cytokeratin (red) in the micro-dissected decidua basalis (**A**: GW 8.2, **B**: GW 8.6), the decidua capsularis (**C**:GW 8.2, **D**: GW 8.6), and the decidua parietalis (**E**: GW 8.2, **F**: GW 8.6). The decidua basalis is characterized by the abundance of trophoblasts cells marked by cytokeratin proteins (CK). Note that glandular epithelial cells are also immunoreactive with anti-CK, whose localization however is different from trophoblast cells. The abundance of of trophoblasts (marker cytokeratin proteins) confirms the histological signatures of decidua basalis.

**Supplementary Figure 3.** Cell-type-specific epigenomic activation of marker genes for each cell type. Cell-type-specific ATAC-seq peaks are often localized in promoter regions of genes marking corresponding cell types. Only selected marker genes are shown here for illustrative purpose. 10Kb upstream regions of each transcription start sites were included. Signal intensities were normalized across all cells.

**Supplementary Figure 4.** When both male and female fetuses were included in the study, the identification of maternal and fetal origins of placental cells can be determined by the abundance of a Y-chromosome gene, *UTY*. Cell types with significant *UTY* expression can be identified as fetal cells. **(A)** The expression of *UTY* on the projected UMAP embeddings across 191,735 detected cells, where fetal cells tended to have increased *UTY* expression. **(B)** Testing on all samples with male fetuses, we observed that the identified fetal cells based on the haplotype analysis by the Souporcell program were highly consistent with the fetal Y-chromosomal gene *UTY* expression, where 94.3% of *UTY*^+^ cells were identified as fetal cells by Souporcell. However, Souporcell, by comparing SNPs between maternal and fetal genomes, has additional power to identify fetal cells when no sequenced reads were mapped onto the *UTY* loci.

**Supplementary Figure 5.** Concordance in cell type annotations between the epigenome (ATAC-seq) and transcriptome (RNA-seq) spaces. Gene activity scores (ATAC-seq) and gene expression were used to identify marker genes for each cell type in the epigenome and transcriptome spaces, respectively, and the percentage of overlapped marker genes for each cell type was used to quantify the concordance between the two modality spaces.

**Supplementary Figure 6.** ChromVar computed regulatory activities of the top transcriptional factors (TF) specific to major cell types. Shown are the top two TFs with the greatest chromVar activity (after normalization) specific to each major cell type.

**Supplementary Figure 7.** *S100A9* displayed increased expression in SCT compared with other trophoblast cells.

**Supplementary Figure 8**. Two clusters of ATAC-seq peaks were identified in the 30Kb upstream region of the *HLA-G* locus, in addition to its promoter ATAC-seq peak. These peaks were specific to EVT cells.

**Supplementary Figure 9.** Enriched expression of *CGA* (**A**), *TFPI2* (**B**), *PLAC* (**C**), and *CCNH* (**D**) in syncytiotrophoblasts (SCT). (**E**) Their enriched gene expression in SCT can also been seen in the Dotplot.

**Supplementary Figure 10.** Expression of the proliferative marker *MKI67* on the UMAP projection.

**Supplementary Figure 11.** Immunolocalization experiments confirmed that PDE3A specifically marks arterial endothelial cells (aECs) in spiral arteries (SA). aECs in the SA with (**A1-A3**) or without trophoblast plugs (**B1-B3**) were immune-reactive to anti-VIM (**A2, B2**), anti-CD31 (**A2, B2**) and anti-PEDE3A (**A3, B3**). Trophoblast cells were marked by cytokeratin proteins (CK). The presence of trophoblast plugs (**A1-A3**) as well as the partial loss of aECs along the vessel wall (**B1-B3**) confirmed the arterial identity of the blood vessels. (**C1-C3**). The absence of PDE3A in venous endothelial cells. The vein in the decidua was not densely surrounded by trophoblast cells (**C1**). The venous endothelial cells (vECs) were immune-reactive to CD31 and VIM, but were negative for the arterial endothelium marker PDE3A (**C3**).

**Supplementary Figure 12.** CODEX profiling on a basal plate sample (GW22) by integrating 10 antibody channels for specific cell types at the maternal fetal interface.

**Supplementary Figure 13. (A-B)** A zoom in view of the CODEX data from a basal plate sample (GW15) showed that villous cytotrophoblasts (VCTs) (arrow indicated) were immunoreactive with anti-CDH1. **(C)** A zoom in view of two SA types from the CODEX data for this GW15 basal plate sample. In contrast to SA-A, endothelial cells lining SA-B was overall negative for PDE3A. However, close examination identified individual PDE3A^+^ endothelial cells along SA-B, suggesting a state transition leading to a loss of PDE3A expression for most endothelial cells in SA-B given venous endothelium cells do not express PDE3A. * denotes the vessel lumen of SA-B.

**Supplementary Figure 14.** A two-channel view of CODEX data displayed co-localization of EVT (marked by CK18) and T cells (marked by CD3) adjacent to a spiral artery.

**Supplementary Figure 15. (A)** 3,514 maternal vascular endothelial cells were profiled in this study, which formed one cluster for arterial endothelial cells (aEC), one cluster for capillary endothelial cells (cEC) and two clusters for venous endothelial subtypes (vEC). **(B)** The known vEC marker *IGFBP7* displayed expected gene expression enrichment in vECs. **(C)** *PDE3A* expression was specific to aECs. **(D)** Representative marker genes identified from our study for maternal vascular endothelial cell subtypes at the maternal-fetal interface. (**E**) The arterial endothelial state transition from the canonical state to the SA-A or SA-B type down-regulated the genes displaying expression enrichment in canonical aECs. Statistical significance was determined by Wilcoxon rank-sum test. *** denoted P < 1e-3 and **** denoted P<1e-4.

**Supplementary Figure 16**. Fate determination of cytotrophoblasts in transient states in anchoring villi. (**A-B**) A group of cytotrophoblast cells were identified in anchoring villi due to their transient states, and these cells can be divided into two groups based on their respective expression specificities of *CD81* (**A**) and *MYCNUT* (**B**) molecules. **(C-F)** The *CD81*^+^ cells have increased SCT marker gene expression exemplified by *CGA* (**C**) and *CYP19A1* (**D**). The MYCNUT^+^ cells have increased EVT marker gene expression demonstrated by *ITGA1* (**E**) and *ITGA5* (**F**). (**G-H**) The *CD81*^+^ cells were more likely to be differentiated into SCT-B cells (marked by GPC5, SCT_GPC5+, Y-axis, **G**) compared with *MYCNUT*^+^ cells, which tended to have a cell fate towards the EVT cell type. The probability of the cell fate commitment was computed and normalized by CellRank. All comparisons were statistically significant with P<1e-4. P values were calculated by Wilcoxon Rank-sum test.

**Supplementary Figure 17.** Different EVT subtypes are marked by different genes. Note that the widely used pan-EVT markers such *as HLA-G*, *ITGA1* and *ITGA5* in fact had expression specificities towards different EVT subtypes.

**Supplementary Figure 18**. UMAP projection of EVT subtypes revealed transcriptomic resemblance between pEVTs and perivascular fibroblasts.

**Supplementary Figure 19.** Expression specificities of key decidual stromal cell marker genes in the five decidual stromal cell subtypes.

**Supplementary Figure 20.** Immunolocalization experiments targeting specific markers to determine the identity and localization of decidual stromal cell subtypes. **(A)** Undecidualized cells (ACTA2^+^) were dispersed in the basal plate (GW20). **(B)** EVT cells (CK^+^) could also express PEDF in the decidua (arrow indicated). Shown is the area in the vicinity of a blood vessel (BV, GW17). **(C)** Cells in the syncytium in anchoring villi were immuno-stained positive for PEDF (arrow indicated). Decidual stromal cells can be identified by their immunoreactivity with anti-VIM and their localization on the maternal side. **(D)** The serial section confirmed that the PEDF^+^ cells in the syncytium were cytotrophoblast cells (CK^+^). The decidual stromal cells immediately adjacent to cell columns were immunoreactive with anti-IGFBP1. **(E-F**) The single FITC (**E**) and merged channels (**F**) showing that the majority of CK^+^ EVTs in the basal plate are Prolactin^+^. (**G, H**) The single FITC (**G**) and merged channels (**H**) showing that the majority of CK^+^ EVTs beneath the anchoring villi are Prolactin^+^.

**Supplementary Figure 21.** Independent confirmation on the specific localization of the DSC4 subtype adjacent to anchoring villi, characterized by their absence of PEDF expression (**A**), and strong expression of IGFBP1 (**A-D**) and PRL (**B-D**).

**Supplementary Figure 22.** Pruning of the DSC3 subtype (DSC3.1) after GW22. The DSC3.1 cells (the right panel) displayed substantially reduced gene expression than the DSC3.2 cells in all main functional categories (the left panel), including anti-angiogenesis, complement activation, antigen presenting and anti-apoptosis.

**Supplementary Figure 23.** A replication study by performing single cell RNA-seq on the decidual swap samples from term labor. **(A)** DSCs were identified from 28,626 sequenced cells by their expression of *IGFBP1*. **(B)** IGFBP1 expression confirmed their decidualization status. **(C)** The presence of *SERPINF1* (PEDF) separated the DSCs into two sub-groups (Path A and Path B). (**D-E**) The subgroup with reduced *SERPINF1* (Path B) displayed increased expression of WNT5A (**D**) and SEMA3A (**E**), a signature also seen from the DSC4 cells in this study.

**Supplementary Figure 24.** The identified endometrial epithelial cells (arrow indicated) in this study had enriched expression of POU5F1 (OCT4, A) and LGR5 (B).

**Legends for Supplementary Tables**

**Table S1.** A summary of existing methods employing single cell approaches to studying placenta (only recent studies were included using high-throughput platforms).

**Table S2.** Summary of sample information, and information for single cell data QC and post-QC statistics (both epigenomic and transcriptome levels).

**Table S3.** Marker genes to identify high-level cell types.

**Table S4.** Cell-type-specific *cis*-acting enhancers in each cell type.

**Table S5.** Antibody information for the CODEX experiments.

**Table S6.** Differentially expressed genes in SA-A and SA-B type aECs compared with canonical ECs. Differentially expressed genes between arterial ECs and venous ECs identified in this study were also included.

**Table S7.** Enriched gene ontology functional terms for the differentially expressed genes in SA-B type aECs compared with canonical ECs.

**Table S8.** Differentially expressed genes and signature genes in EVT subtypes.

**Table S9.** Enriched gene ontology and MGI functional terms for genes specific to EVT subtypes.

**Table S10.** Differentially expressed genes between Path A and Path B, and among terminal cell states.

**Table S11.** Enriched gene ontology functional terms for genes with specific expression in Path A and Path B.

**Table S12**. Communication probabilities between arterial endothelial cells (aECs) and DSCs recovered by CellChat across 27 signaling pathways.

**Table S13.** Disease relevance scores associated with pre-eclampsia (PE), spontaneous preterm labor (sPTB) or miscarriage across cell types.

**Table S14.** Information for all antibodies used in immunochemical experiments in this study.
